# Conservation of an Ancient Oxidation Response That Controls Archaeal Epigenetic Traits Through Chromatin Protein Networks

**DOI:** 10.1101/801316

**Authors:** Sophie Payne, Marc Facciotti, Kevin Van Cott, Andrew Yao, Mark Wilson, Stephan Sutter, Kiara L. Rodríguez-Acevedo, Paul Blum

## Abstract

Epigenetic variants of the archaeon *Sulfolobus solfataricus* called SARC have evolved heritable traits including extreme acid resistance, enhanced genome integrity and a conserved “SARC” transcriptome related to acid resistance. These traits appear to result from altered chromatin protein function related to the heritable hypomethylation of chromatin proteins Cren7 and Sso7D. To clarify how this might occur, ChIPseq and Affinity Purification Mass Spectrometry (AP-MS) were used to compare Cren7 and Sso7D genome binding sites and protein networks between lineages (wild type and SARC) and culture pH (pH 1 and 3). All SARC transcriptome loci were bound by these chromatin proteins but with invariant patterns indicating binding alone was insufficient to mediate the SARC traits. In contrast, chromosome association varied at other loci. Quantitative AP-MS was then used to identify protein interaction networks and these included transcription and DNA repair proteins implicated in the evolved heritable traits that varied in abundance between SARC and wild type strains. Protein networks included most of the S-adenosylmethionine (SAM) synthesis pathway including serine hydroxymethyltransferase (SHMT), whose abundance varied widely with culture pH. Because epigenetic marks are coupled to SAM pools and oxidative stress in eukaryotes, occurrence of a similar process was investigated here. Archaeal SAM pools were depleted by treatment with SAM pathway inhibitors, acid or oxidative stress and, like eukaryotes, levels were raised by vitamin B12 and methionine supplementation. We propose that in archaea, oxidation-induced SAM pool depletion acting through an SHMT sensor, drove chromatin protein hypomethylation and thereby protein network changes that established the evolved SARC epigenetic traits.

**Significance Statement:** Archaea and eukaryotes share many molecular processes, including chromatin-mediated epigenetic inheritance of traits. As with eukaryotes, archaeal protein complexes were formed between trait-related proteins and chromatin proteins, subject to chromatin protein methylation state. Oxidation-induced depletion of S-adenosylmethionine (SAM) pools likely resulted in chromatin protein hypomethylation. Subsequent chromatin enrichment of serine hydroxymethyltransferase as a response to oxidative stress could modulate methylation at specific genomic loci. The interplay between archaeal metabolism and chromatin appear consistent with patterns observed in eukaryotes and indicate the existence of an ancient oxidation signal transduction pathway controlling epigenetics.

## Introduction

Epigenetics is defined as the heritability of a phenotypic state that does not result from changes in DNA sequence. In eukaryotes, epigenetic mechanisms involve reversible post-translational modification of histones or DNA and are essential for influencing adaptive traits and cellular development. Components of chromatin-mediated epigenetics require histones, histone modification writers, readers and erasers (1), and histone remodelers (2). DNA methylation requires enzymes that deposit, read and erase CpG methylations (3). Together these mechanisms underly epigenetic processes across eukaryotic lineages (4).

The domain archaea share molecular processes with eukaryotes using homologous proteins to accomplish them. These include proteins involved in DNA replication, DNA repair, RNA transcription and protein translation (5,6). Due to these conserved features and more recent phylogenetic work, a two domain tree model was proposed in which eukaryotes arose from archaea (7). Until recently however, it was unclear if epigenetics was included among these processes. Although the euryarchaeotal phylum use histone homologs (8) to package DNA, they are not post translationally modified (9) and lack epigenetic activity. However, recent work indicates epigenetic mechanisms are operative in the crenarchaeotal phylum (4,10–12).

The crenarchaeote and thermoacidophile, *Sulfolobus solfataricus*, was used in adaptive laboratory evolution experiments to improve acid resistance (11). Surprisingly, this and additional evolved traits appeared to arise through an epigenetic mechanism (4,10,12). Triplicate trials involving serial passage at sequentially lower pH values produced independent cell lines named super-acid-resistant Crenarchaeota (SARC) (11). All SARC had a 100-fold increase in acidophily and 10-fold increase in resistance to the common oxidative microbicide sulfite, supporting the link between pH and oxidative stress (11–14). These traits were accompanied by a conserved and heritable transcriptome of “SARC genes” related to acid tolerance that was retained after passage without selection (11,12). In addition, SARC had increased genome integrity manifested as reduced mutation and transposition rates relative to a control strain passaged without selection. Although mutations that explained the SARC traits were predicted, the genome of one of the SARC lines (SARC-I), had no mutations, transpositions or rearrangements relative to its parent (SULG) after extensive PCR, resequencing and bioinformatic analysis (12). Therefore, an epigenetic mechanism was proposed to explain the evolved traits (12). Moreover, it was found that gene expression and phenotype could be perturbed in SARC using recombination to replace the native chromatin at SARC gene loci with naïve DNA, without introducing sequence change (12). As DNA methylation in archaea is bacterial-like, the eukaryotic epigenetic mechanism of CpG site methylation is unlikely to occur in *Sulfolobus* (15). However, because chromatin proteins Sso7D (16) and Cren7 (17) are heritably hypomethylated (10), the SARC traits may instead be mediated by Cren7 and Sso7D and their modification state in a process analogous to that of undermodified eukaryotic histones.

In eukaryotes, a link has been established between epigenetics, oxidative stress and metabolism via the SAM and one-carbon pathways, that produce substrate for histone methylation (18,19). Reactive oxygen species inactivate the SAM-generating enzyme methionine synthase (20) and its cofactor vitamin B12 (21). Without a methyl donor, histones become undermethylated (22) and expression patterns may be altered via perturbed protein-protein interactions (23). To maintain certain histone methylations even during oxidative SAM limitation, the eukaryotic SAM synthase II isozyme directly associates with a histone and histone methyltransferase to promote methylation (24). In addition, the eukaryotic SESAME complex contains SAM pathway enzymes and interacts with the histone methyltransferase Set1 (25). New evidence in eukaryotes suggest that histones store methyl groups and that histone undermethylation is a controlled process (26). In *S. solfataricus*, the calculated protein-bound methyl pool (27) is two orders of magnitude greater than the soluble SAM pool per cell (28), supporting this idea. The studies described here were undertaken to identify factors controlling archaeal chromatin protein methylation that might link epigenetics, oxidative stress and metabolism.

## Methods

### Archaeal strains and cultivation

*S. solfataricus* (SULG) (12,29) and an acid-adapted derivative of this cell line (SARC-I) (12) were grown as described previously by Payne et al. (12) and SI Appendix, Supplemental Methods. Cells were harvested at an OD_540nm_ of 0.5, were crosslinked as described previously (30) and stored as 20OD pellets at −20C until further use. Media reductive potential was measured using a Pinpoint Platinum ORP/redox Probe (American Marine).

### Immunoprecipitation for ChIP-Seq

Triplicate crosslinked cell pellets were sonicated and subjected to chromatin immunoprecipitation using polyclonal antibody serum for Cren7 or Sso7D as described previously (30) with alterations described in and SI Appendix, Supplemental Methods.

### ChIP-Seq and data analysis

DNA samples were blunt ended, A-tailed, ligated to barcoded adapters, PCR amplified and sequenced as described previously (30) and in SI Appendix, Supplemental Methods. ChIP and WCE DNA libraries were demultiplexed, processed and analyzed as described in (30) and SI Appendix, Supplemental Methods.

### Immunoprecipitation for AP-MS

Duplicate crosslinked pellets were sonicated and immunoprecipitated, with DNAse treatment as described in SI Appendix, Supplemental Methods.

### LC-MS/MS of ChIP samples

Protein samples were digested, purified and subjected to mass spectrometry for identification and quantitation relative to a spiked standard as described in SI Appendix, Supplemental Methods.

### Quantitative AP-MS data analysis

The MS/MS spectra were searched against the *Sulfolobus solfataricus* proteome and immunoprecipitated protein abundances were compared to samples that controlled for background, as described in SI Appendix, Supplemental Methods.

### SAM pool measurement

SAM was extracted by solid phase extraction as in Struys, et al. (31), with modifications as described in SI Appendix, Supplemental Methods.

### Data availability

ChIPseq data are available in the Gene Expression Omnibus archive, with accession numbers in the SI Appendix, Supplemental Methods.

## Results

### Genome distribution of chromatin proteins

Prior work indicated that heritable hypomethylation of chromatin proteins and their binding to particular genes may mediate the SARC traits (10,12). However, these proteins bind the minor groove and their methylated residues are on solvent-facing surfaces (10) that do not appear to alter DNA-binding affinity (17,32). Therefore, it was predicted that the location and binding affinity of these proteins was invariant between wild type and SARC strains. To test this predictions, ChIPseq was conducted using wild type and SARC (12) strains to determine Cren7 and Sso7D genome binding patterns. In this work, parental SULG (henceforth referred to as wild type) and its evolved epigenetic variant SARC-I (SARC) were used because they are genetically identical but phenotypically distinct (12).

Cren7 and Sso7D proteins exhibited genome-wide coverage (Fig. 1; SI Appendix Fig S1&S2) and bound all SARC transcriptome genes including those whose expression was perturbed by recombination (12), supporting their role in the SARC traits. ChIPseq peaks often overlapped between the two proteins. As predicted, the genome location and read depth of most Cren7 and Sso7D peaks did not vary greatly between wild type or SARC strains cultured under identical conditions (Fig. 1). In addition, the size of peaks and their change in size between wild type and SARC cultured under identical conditions did not correlate with patterns of expression level or expression change in SARC (12). Therefore, while chromatin protein presence may contribute to chromatin-mediated transcriptional regulation, DNA-binding affinity does not. Peak patterns at SARC genes were consistent between conditions, while peak variation occurred at low frequencies for other loci. Both proteins had high affinity (high sequence read depth) (SI Appendix Table S1&S2) and low affinity (low sequence read depth) peak patterns in which Cren7 binding peaks were fewer and larger relative to Sso7D, while Sso7D exhibited more and smaller peaks. This appeared consistent with Cren7’s greater binding affinity compared to Sso7D (17,33). However, both proteins had a similar genome coverage of 45%. This is because the larger width of Cren7 peaks provided a similar genome coverage as many but narrower low affinity Sso7D peaks. These data suggest that Cren7 is more targeted in its interaction with the genome while Sso7D is more general. Although peak patterns were similar between strain type, the size of most high affinity Sso7D peaks in SARC increased when cultured at lower pH (pH 1) (Fig. 1, SI Appendix Fig S1&S2).

**Figure 1.**
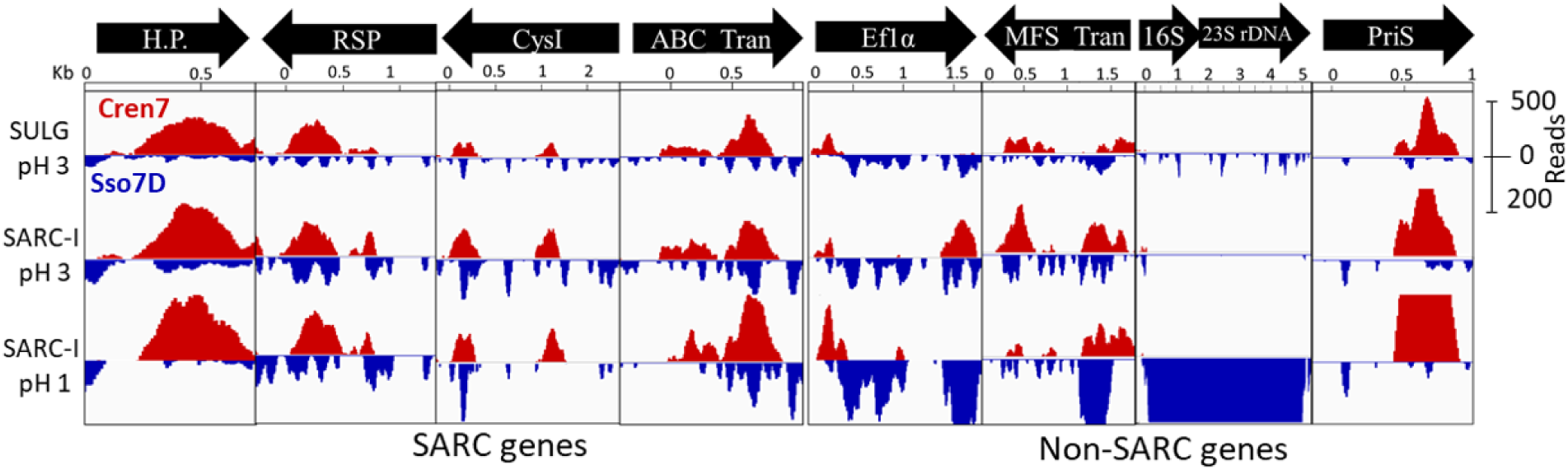
Representative genome binding patterns of Cren7 and Sso7D in the *S. solfataricus* genome using ChIPseq. Y-axis indicates ChIP-seq read depth, bars represent read depth averaged over 10bp (red = Cren7 ChIP, blue = Sso7D ChIP). The maximum displayed peak height is 500 reads for Cren7 and 200 reads for Sso7D. Genes: H.P = Hypothetical protein SULA 0759, RSP = Radical SAM domain protein SULA 0674, CysI = Sulfite reductase SULA 0698, ABC Tran = Peptide ABC transporter SULA 2263, ef1α = elongation factor 1 alpha SULA 0225, MFS Tran = MFS transporter SULA 0327, 16S/23S rRNA = rRNA loci, priS = DNA primase SULA 2052.

The ORFs associated with high affinity binding sites for both chromatin proteins appeared related to SARC traits (SI Appendix Table S1&S2). Cren7 had 157 high affinity sites that included ORFs involved in SAM metabolism, DNA repair, oxidation resistance, transcriptional regulators, CRISPR Cas enzymes and several transposases (SI Appendix Table S1). Sso7D had 64 high affinity sites that included ORFs present in the Cren7 dataset, with the addition of ORFs involved in RNA processing, DNA remodeling and many IS4 family transposases (SI Appendix Table S2). Each protein also exhibited unique gene associations. Cren7 was highly abundant at CRISPR array loci amounting to 47% of its total genome binding and may form a chromatin fiber (34). In contrast, Sso7D was abundant at the rDNA locus, but only in SARC at pH 1, amounting to 4% of its total genome binding while Cren7 was absent at this locus (Fig. 1). Sso7D was also bound at many insertion sites of insertion sequence element IS4. This is intriguing because the “cut and paste” IS4 element was the most active IS family in SUL120, a control cell line passaged without acid selection whose mutation and transposition rates were higher than in SARC (12). In contrast, Cren7-bound transposase genes were not active in SUL120. These results suggest Cren7 may be important in immunity to foreign DNA, while Sso7D may be important for protein synthesis and may inhibit IS element excision in the cut and paste transposition mechanism. Furthermore, the broad distribution of Sso7D compared to Cren7 may indicate a role in DNA protection such as suggested for bacterial Dps (35). Again, these binding pattern differences suggest the two proteins fulfill different biological roles that may facilitate the SARC traits.

The consistency of peak location between strain and culture condition implied a non-random DNA binding mechanism. Despite this, the high affinity peaks for both proteins were not bound at conserved DNA sequences or regions of GC bias, which is consistent with previously reported data (17,32). Searches for potential binding motifs limited motif size to at least 6bp as this is the smallest observed binding site of both proteins (17,33) and avoids statistically frequent smaller motifs. Therefore, since both proteins bind the minor groove and have limited capacity to bind specific sequences (16,17), their non-random binding sites must arise from their association with other DNA binding proteins. A similar mechanism has been observed with *Sulfolobus* SS-LrpB, which binds DNA when associated with LysM (36).

### Archaeal chromatin protein networks

In eukaryotic histones, methylation changes on the solvent facing surfaces of histone cores do not alter DNA-binding affinity but can dramatically alter transcription and DNA repair through protein-protein interactions (23,37). Johnson et al found Cren7 and Sso7D were modified at solvent facing residues and not at DNA-contacting sites (10). Furthermore, methylation state does not alter DNA-binding affinity (17,32). Because SARC exhibited traits associated with transcriptional regulation and the preservation of genome integrity (12), it was predicted that Cren7 and Sso7D associate with other proteins involved in DNA binding, transcription, and DNA repair. Since ChIPseq patterns differed between the two chromatin proteins, protein complexes also were likely to differ in a manner consistent with different biological function. Furthermore, chromatin protein undermethylation in SARC was predicted to alter protein-protein association affinities. To test these predictions, quantitative affinity purification and mass spectrometry (AP-MS) (38) of crosslinked chromatin protein complexes was used to identify and quantify the abundance of proteins present in the protein interaction network (interactome) of Cren7 and Sso7D in wild type cultured at pH 3 and SARC cultured at pH 3 and 1. Comparison to the existing transcriptomic dataset (12) permitted an integrated analysis of differential protein interactions that accounts for changes in expression level. It should be noted that AP-MS cannot distinguish between interactions that are direct or indirect but do indicate participation in protein complexes. This method is quantitative through the addition of an internal standard, allowing the determination of a protein’s relative change in abundance between strains and pH conditions.

As predicted, the interactome components for both chromatin proteins often differed in abundance for wild type and SARC cultured under identical conditions (Fig. 2, SI Appendix Table S3&S4). These changes were not explained by transcriptional changes at the cognate genes (12). Because the wild type and SARC strains have identical genome sequences, these differences could result from an epigenetic process. Proteins in the interactome of Cren7 and Sso7D generally occurred at similar abundances for wild type cultured at pH 3 and SARC cultured at pH 1 (Fig. 2). However, when cultured at the same pH, the total abundance of proteins associating with Cren7 in SARC was 12% ± 2 less compared to the parental strain. For Sso7D, the total protein abundance was 60% ± 10 more in SARC. Many proteins were involved in functions related to the SARC traits, including transcription, chromosomal topology and repair, translational regulation, and the SAM synthesis pathway (Fig. 2, SI Appendix Table S3&S4). As predicted from overlapping ChIPseq patterns, Sso7d appeared in the Cren7 interactome. Method sensitivity may explain the absence of Cren7 in Sso7D’s interactome. The Cren7 interactome included key components of the basal transcription complex such as TATA-box binding protein (TBP), transcription initiation factor IIB (TFB), transcription factor S (TFS) and the Rpo3, 4, 7 and 11 subunits of RNA polymerase, supporting a role in transcriptional regulation. The Cren7 interactome also contained a putative Ruv-B protein, aTIP49 protein and a Type II topoisomerase VI, which are important for recombination and DNA repair (39,40).

**Figure 2.**
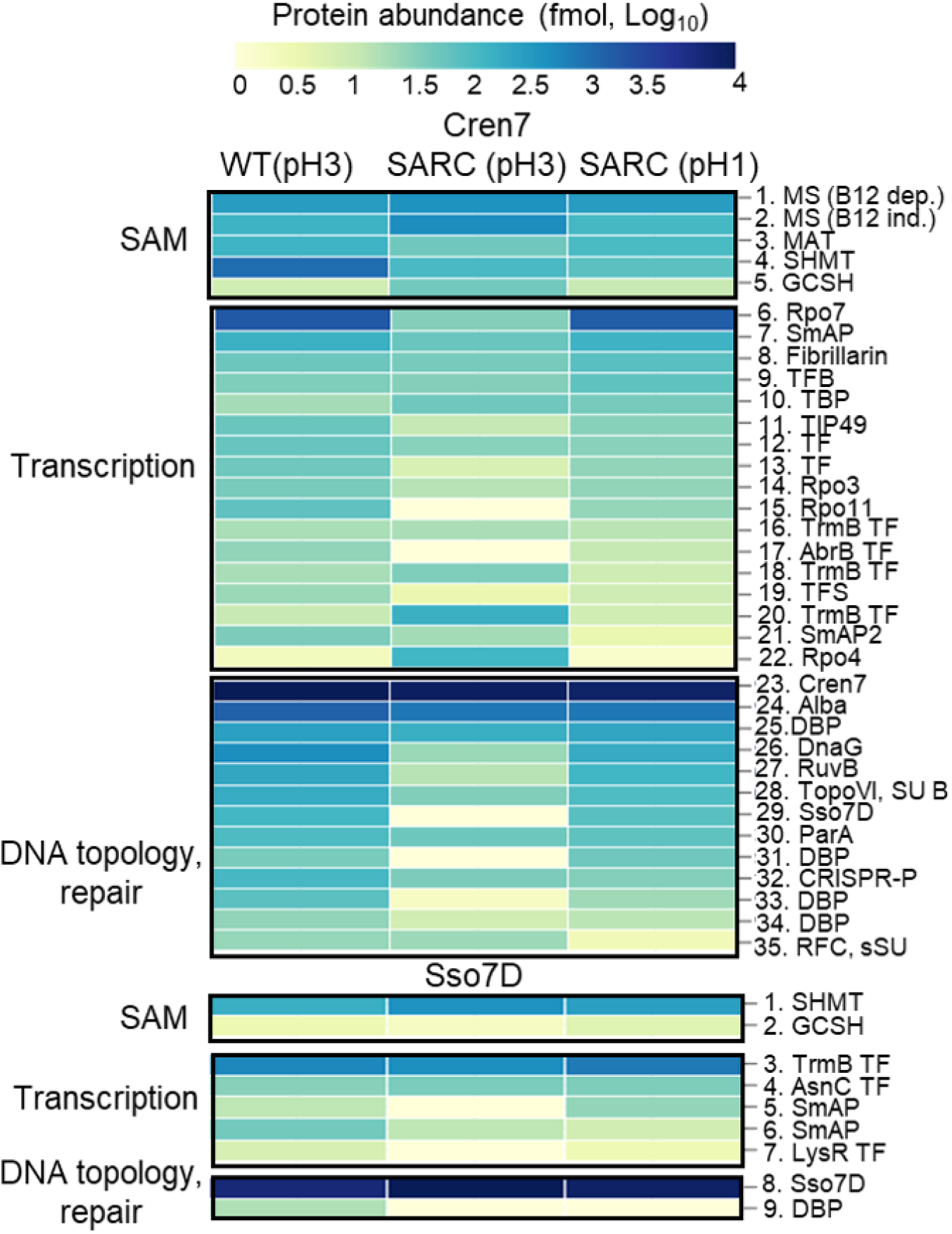
Abundance heatplot of select proteins in Cren7 and Sso7D interactomes. Color indicates protein abundance in fmol per 5×109 cells on a Log10 scale. Protein reference numbers refer to SI Tables 3 & 4.

Interestingly, the Cren7 interactome also included most proteins in the SAM synthesis pathway including SAM synthase, B12-dependent and independent methionine synthases (MS), serine hydroxymethyltransferase (SHMT) and glycine cleavage system H protein (GCSH). Sso7d also interacted with SHMT and GCSH. Only some proteins made contacts with both Sso7D and Cren7 interactomes. Both proteins interactomes included various transcriptional regulators (TFs) and other DNA-binding proteins, which are likely to confer chromatin protein sequence specificity and may alter regulatory activity related to SARC traits (36,41,42). Additional interactome features included protein abundance differences when comparing wild type and SARC and comparing SARC growth pH conditions (Fig. 2, abundance values and sample variance in SI Appendix Table S3&S4). For the Cren7 interactome, proteins related to SAM metabolism may be involved in regulating chromatin protein methylation state. SAM synthase was decreased −2-fold for SARC at pH 3 relative to wild type at pH 3, while MS and GCSH were increased (4 and 4.5-fold). SHMT was highly increased (9.4-fold) for SARC at pH 1 relative to the two other conditions. Proteins related to transcription were also altered and may be involved in the heritable transcriptional changes in SARC. Proteins that were decreased more for SARC at pH 3 relative to wild type at pH 3 included Rpo3, Rpo7, Rpo11 (−2, −50-fold, and absent), TFS (−2.6-fold), TFB (−2-fold), and two TFs (−4.5-fold and absent), while two TFs and Rpo4 were increased (4.3, 18.8 and 93-fold, respectively). TBP was decreased −2.7-fold for SARC at pH 1 relative to SARC at pH 3.

The abundance of Cren7 interactome proteins related to DNA topology and repair also changed and may be involved in the reduced mutation rates of SARC. Proteins that decreased more for SARC at pH 3 relative to wild type at pH 3 included a topoisomerase (−2.7-fold), Rub-B like protein (−9.1-fold), a DNA binding protein (−12.5-fold) and Sso7D (absent). Proteins that were increased for SARC at pH 1 relative to SARC at pH 3 included two DNA binding proteins (41.2 and 3.5-fold), DNA primase (19.7-fold), a putative RuvB (14.8-fold) and a CRISPR-associated protein (2.9-fold). In the Sso7D interactome, a LysR TF related to transcription was increased 2.1-fold for SARC at pH 3 relative to wild type. For DNA topology and repair, a DNA binding protein was associated for SARC at pH 1 but not for other conditions.

### SAM pools and oxidative stress

In eukaryotes, maintenance of chromatin histone methylation state depends on S-adenosyl methionine (SAM) pool size and is affected by oxidative stress. This is because oxidative stress inactivates methionine synthase and its cofactor vitamin B12 that are required for SAM synthesis (18,22). Oxidative stress is a relevant stress for *S. solfataricus* because it occurs during cultivation at high temperature and low pH. *Sulfolobus* medium measured for oxidation-reduction potential (E_h_), a measure of oxidative stress (43), shows that it is a more oxidizing environment at pH 1.0 (E_h_ 648mV) than pH 3.0 (E_h_ 504mV). Therefore, it was predicted that SAM pools would be depleted in SARC when cultured at higher E_h_ (lower pH) values. If true, then a metabolic basis for hypomethylation could be established. To test this hypothesis, SAM was extracted from wild type and SARC strains under various growth conditions and measured using HPLC.

The two strains had similar SAM pools and were similarly affected by treatment conditions (Fig.3 for SARC, SI Appendix Fig.3 for wild type). SAM pools were depleted by exposure to low pH (high E_h_), as they were 43% lower for SARC grown continuously at pH 1.0/E_h_ 648mV (110 µM SAM ± 2) than for SARC grown at pH 3.0/E_h_ 504mV (192µM SAM ± 8). As SAM depletion leads to hypomethylation of eukaryotic histones, the same likely occurred with SARC chromatin proteins (18,22). This depletion occurred rapidly, as cells transferred from continuous pH 3 culture to pH 1 for 1 hr had SAM pools reduced by 22% (150µM SAM ± 4). Furthermore, oxidative stress in the absence of acid stress also depleted SAM, further supporting the connection between pH, oxidative stress and SAM metabolism (13,14,19,22). For example, SAM pools in cells cultured at pH 3 and treated with 120µM hydrogen peroxide for 1 hr were 71µM ± 4, a 63% reduction compared to untreated cultures (Fig. 3). The concentration of SAM in wild-type *S. solfataricus* determined here was 162 µM ± 6 which is close to previously reported values (28) and to those of eukaryotes (44).

**Figure 3.**
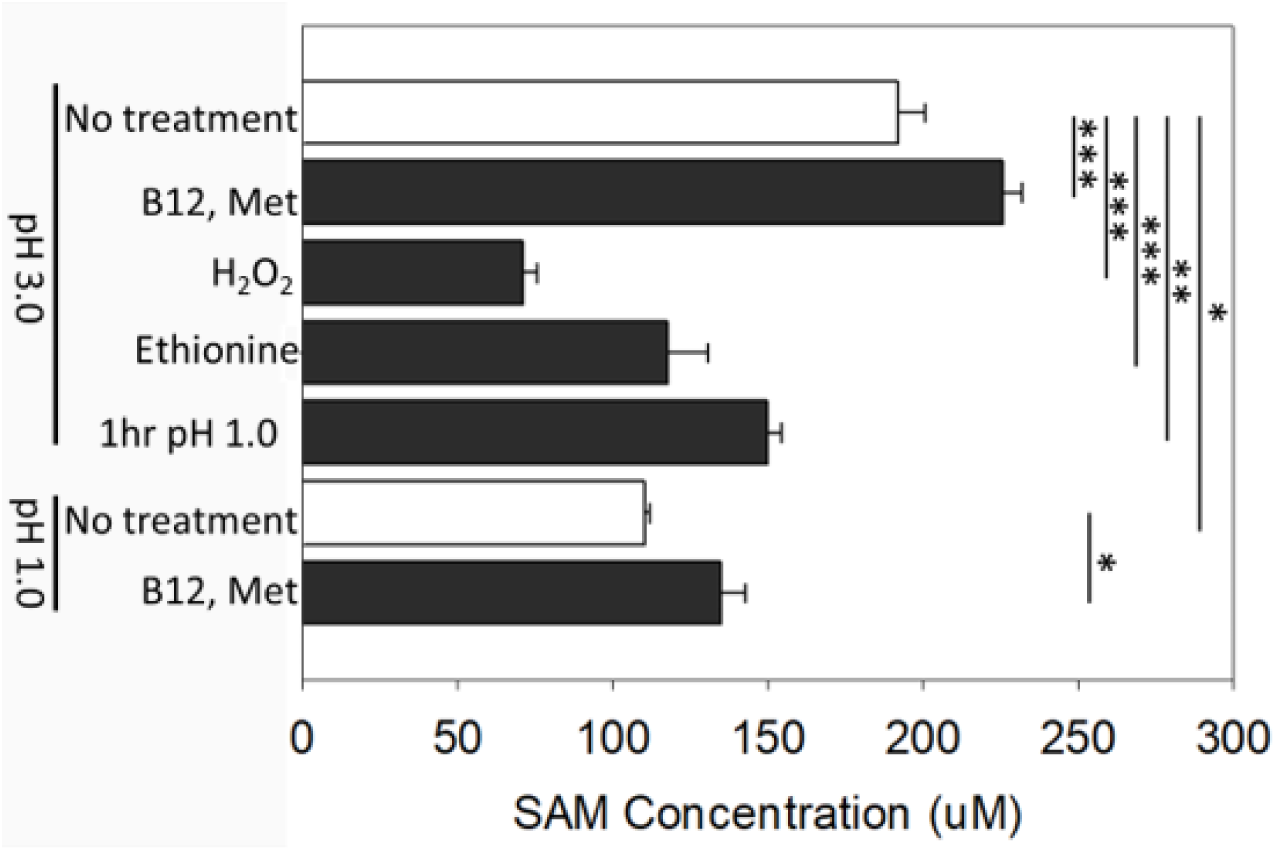
Effect of treatments on S-adenosylmethionine (SAM) abundance in SARC. SARC-I was passaged 3x at pH 3.0 (Eh 504mV) or pH 1.0 (Eh 648mV). pH 3.0 cultures were treated with oxidative stress (120uM H2O2) for 1hr prior to SAM extraction, with 1mM ethionine during culture, with 500nM Vitamin B12 and 10mM methionine for 12hrs prior to SAM extraction, or transferred to pH 1.0 media for 1hr prior to extraction. pH 1.0 cultures were also treated with 500nM Vitamin B12 and 10mM methionine for 12hrs prior to SAM extraction. n = 4 for all conditions. (*) P < 0.05, (**) P < 0.01, (***) P < 0.001, Students T-test

To determine if SAM depletion at increased E_h_ resulted from inhibition of the B12 and SAM synthesis pathways, the response of SAM pool levels was evaluated by manipulating these pathways. Methionine adenosyltransferase is essential for SAM synthesis and is inhibited by ethionine (45). Cells cultured at pH 3 were treated with 1mM ethionine, which depleted SAM pools by 39% (118µM SAM ± 13). To relieve metabolic bottlenecks, pH 1 and pH 3 cultures were supplemented with 500nM vitamin B12 and 10mM methionine. Supplementation increased SAM by 17% (226µM SAM ± 6) for pH 3 cultures and 23% (135µM SAM ± 8) for pH 1 cultures (Fig. 3). These data indicate SAM pools can be manipulated by oxidative stress and pH, and provide a physiological route to establish epigenetic marks in SARC.

## Discussion

By analogy to eukaryotic histone-mediated epigenetics, hypomethylation of SARC chromatin proteins Cren7 and Sso7D could play a similar role in the epigenetic traits observed in SARC. To test this, the role of chromosomal binding location and affinity was measured using ChIPseq, and protein interaction networks were identified using AP-MS. Although genomic binding patterns of chromatin proteins Cren7 and Sso7D did not vary between wild type and SARC strains under identical conditions, protein interactome networks did. This is consistent with the minor groove binding of these proteins, and that methylated residues are solvent facing and would affect protein-protein interactions but not DNA binding affinity. Interestingly, the Cren7 protein interactome contained much of the SAM synthesis pathway which appeared conserved with the eukaryotic pathway despite the evolutionary distance. This included SHMT, whose abundance increased 9.4-fold under pH 1 culture conditions, contradicting its expression levels (12). Furthermore, SAM pools were depleted under oxidative conditions, a response that is conserved in eukaryotes and was likely experienced by SARC during their evolution. These findings establish a metabolic basis for chromatin protein hypomethylation and are consistent with a model for the role of chromatin methylation state in the SARC traits (Fig. 4).

Cren7 forms network contacts with most of the basal transcription complex, indicating its potential role in regulating gene expression (Fig. 4A). The abundance of these components varies for SARC compared to wild type cultured under identical conditions. This implicates an epigenetic effect on the transcription complex, mediated by the heritable modification state of Cren7. The exact mechanism for this mode of transcriptional regulation is currently unknown. A regulatory link with SAM metabolism is also proposed (Fig.4B). SAM pools became depleted during the evolution of SARC. However, the methylation of certain chromatin regions may confer an essential expression state and must be maintained regardless of SAM abundance. For many pathways, protein components form complexes to allow for substrate tunneling, increasing biochemical efficiency, controlling flux at network branch points, and overcoming substrate limitations (46). The SAM pathway may form a complex that associates with Cren7, channeling substrates and increasing local SAM abundance (Fig. 4B), which is consistent with patterns observed in eukaryotes (24,25). In the Cren7 interactome, the SAM pathway is present for all strains and growth conditions, while SHMT is much more abundant at pH 1. However, SHMT is 9.4-fold more abundant for SARC cultured at pH 1, which is inconsistent with its expression levels (12). SHMT may therefore sense cytosol redox state, associating with the Cren7-complexed SAM pathway to improve Cren7 methylation at certain sites during SAM depletion (Fig. 4B).

**Figure 4.**
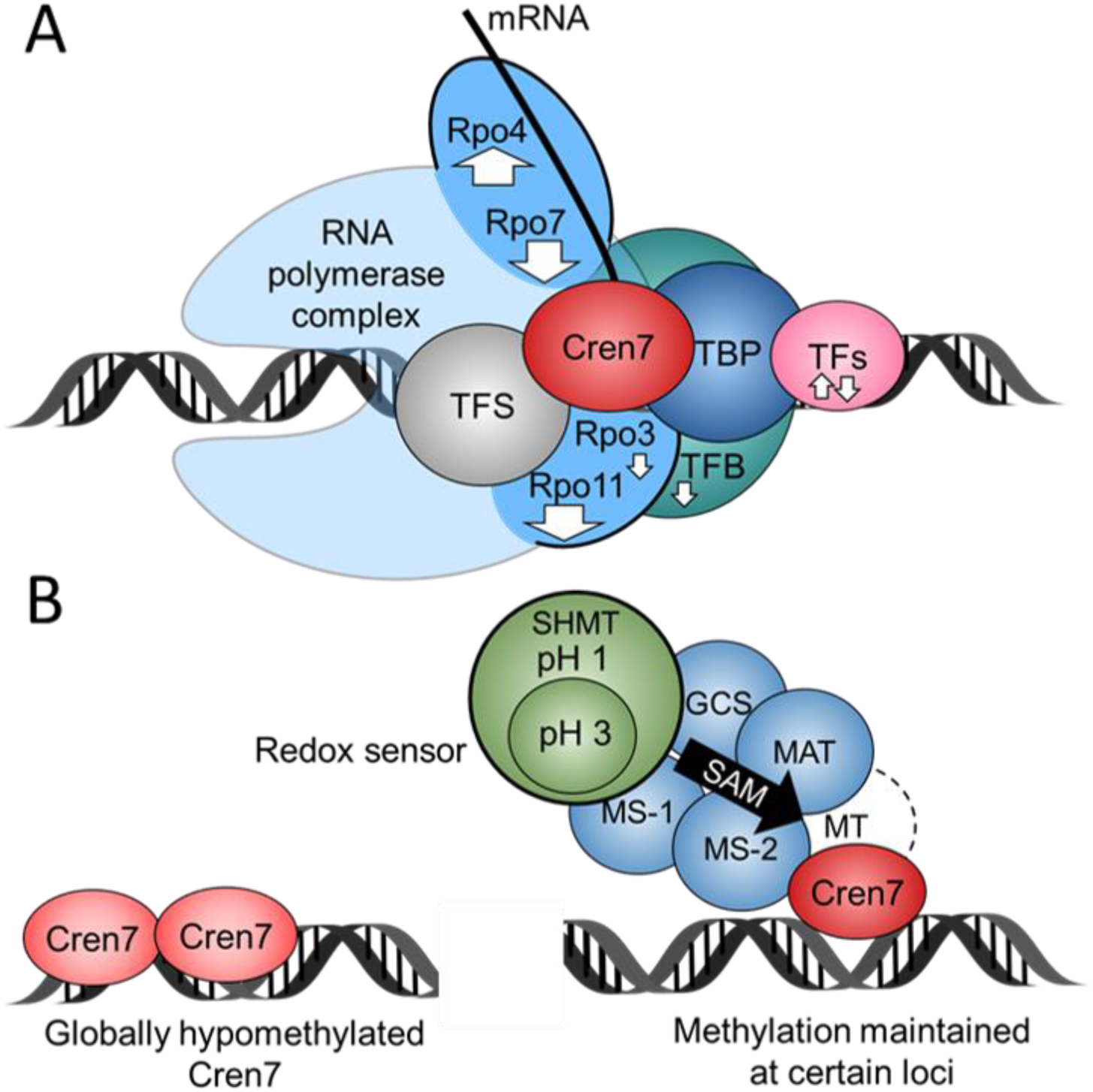
Proposed model for the role of Cren7 interactome proteins in the SARC traits. A) Cren7 interaction with the basal transcription complex. Cren7 in the transcription complex may regulate transcription by recruiting, blocking, or stabilizing other proteins. Interaction affinity is increased (up arrow) or decreased (down arrow) in SARC relative to wild type cultured under identical conditions (arrow size = degree of change). Proteins: TATA-box binding protein (TBP), Transcription initiation factor IIB (TFB), Transcription elongation factor S (TFS), Various transcriptional regulators (TF’s), RNA polymerase subunits (Rpo#). B) SAM metabolic pathway enzymes interacting with Cren7 constitute a redox sensor (SHMT) that regulates chromatin modification. The SAM synthesis pathway complexes with Cren7 to channel SAM and ensure methylation of associated chromatin regions. SHMT abundance increases under oxidative stress, further increasing substrate flow. Enzymes: Methionine synthase (MS1-B12 dependent, MS2 – B12 independent), serine hydroxymethyltransferase (SHMT), methionine adenosyltransferase/SAM synthase (MAT), Glycine cleavage system (GCS), unknown Cren7 methyltransferase (MT)

Because the methylated residues of Cren7 and Sso7D are solvent facing (10), it was predicted that their ChIPseq binding patterns would be present at SARC genes but would not change between wild type and SARC cultured under identical conditions. As expected, all SARC genes were bound by one or both chromatin proteins and those patterns did not vary between strains. This indicates that while chromatin protein presence may contribute to chromatin-mediated transcriptional regulation, DNA-binding affinity does not. Furthermore, the absence of a conserved binding sequence and the minor-groove binding location of Cren7 and Sso7D, suggest that protein binding partners might instead confer binding specificity. As all previous work on Cren7 and Sso7D DNA binding was performed in vitro, the findings presented here provide important insight into the *in vivo* DNA-binding patterns of archaeal chromatin proteins. Structural comparisons indicate that the two proteins differ in how their DNA-binding surfaces interact with DNA (47). As predicted, the two had different binding patterns and exhibited distinctly different dynamics in their responsiveness to genetic background and changing environmental conditions. For example, the large amount of Cren7 bound to the CRISPR arrays may serve as a chromatin protein repository for regulatory changes or for populating naïve DNA during cellular replication.

Because genome binding patterns were unaltered in SARC, it was predicted that Cren7 and Sso7D formed protein interaction networks with proteins involved in SARC traits, and that these interactions were altered in SARC. As expected, the two proteins interacted with SARC trait-related proteins and might regulate these various functions by altering complex composition or protein activity. The Cren7 interactome included more proteins related to SARC traits, which is consistent with the notion that Cren7 is more highly conserved across the phylum *Crenarchaoeta* and is more likely to mediate a conserved mechanism of epigenetics. Interestingly, although a known archaeal MT with broad substrate specificity, aKMT, methylates Cren7 in vitro (48), it was not found in the protein networks presented here. Archaea lack homologs to known eukaryotic histone MTs so additional work may lead to their identification.

Protein networks could result from interaction with both soluble and DNA-bound chromatin protein however, it is thought that at least Cren7 occurs primarily in the DNA-bound state (17). Further work is required to distinguish between these interactions and to identify the chromatin composition of specific chromosomal loci. “Cytosolic” proteins that appear may be interacting with the soluble fraction. However, such proteins are often present in the eukaryotic nucleus and have regulatory effects (49). As archaea are deficient in multi-domain proteins (50) and *Crenarchaeota* are deficient in two-component regulatory systems (51), these metabolic proteins may have regulatory interactions with chromatin and represent signal transduction pathways. Some archaeal proteins have motifs that appear to be proto-nuclear localization signals (52), which may also explain their presence near the chromosome. The physical association between Cren7 and the SAM pathway suggests a connection between 1-carbon metabolism and Cren7 methylation state, providing a mechanism for regulating the occurrence of epigenetic traits. As the interaction between oxidative stress and cellular metabolism is ancient (53), the data presented here indicate that epigenetic systems originally evolved in the context of balancing SAM pools for protein methylation and redox homeostasis (19–21).

## Acknowledgements

This work was supported by the National Science Foundation Grant Molecular and Cellular Biosciences (MCB)-1517408.

## References

1. Hyun, K., Jeon, J., Park, K., and Kim, J. (2017) Writing, erasing and reading histone lysine methylations. Exp Mol Med 49

2. Becker, P. B., and Workman, J. L. (2013) Nucleosome Remodeling and Epigenetics. Csh Perspect Biol 5

3. Klose, R. J., and Bird, A. P. (2006) Genomic DNA methylation: the mark and its mediators. Trends Biochem Sci 31, 89–97

4. Blum, P., and Payne, S. (2019) Evidence of an Epigenetics System in Archaea. Epigenet Insights 12, 2516865719865280

5. Gupta, R. S. (1998) Protein phylogenies and signature sequences: A reappraisal of evolutionary relationships among archaebacteria, eubacteria, and eukaryotes. Microbiol Mol Biol R 62, 1435–+

6. Puhler, G., Leffers, H., Gropp, F., Palm, P., Klenk, H. P., Lottspeich, F., Garrett, R. A., and Zillig, W. (1989) Archaebacterial DNA-Dependent Rna-Polymerases Testify to the Evolution of the Eukaryotic Nuclear Genome. P Natl Acad Sci USA 86, 4569–4573

7. Williams, T. A., Foster, P. G., Cox, C. J., and Embley, T. M. (2013) An archaeal origin of eukaryotes supports only two primary domains of life. Nature 504, 231–236

8. Sandman, K., and Reeve, J. N. (2006) Archaeal histones and the origin of the histone fold. Curr Opin Microbiol 9, 520–525

9. Forbes, A. J., Patrie, S. M., Taylor, G. K., Kim, Y. B., Jiang, L., and Kelleher, N. L. (2004) Targeted analysis and discovery of posttranslational modifications in proteins from methanogenic archaea by top-down MS. Proc Natl Acad Sci U S A 101, 2678–2683

10. Johnson, T., Payne, S., Grove, R., McCarthy, S., Oeltjen, E., Mach, C., Adamec, J., Wilson, M. A., Van Cott, K., and Blum, P. (2019) Methylation deficiency of chromatin proteins is a non-mutational and epigenetic-like trait in evolved lines of the archaeon Sulfolobus solfataricus. J Biol Chem 294, 7821–7832

11. McCarthy, S., Johnson, T., Pavlik, B. J., Payne, S., Schackwitz, W., Martin, J., Lipzen, A., Keffeler, E., and Blum, P. (2016) Expanding the Limits of Thermoacidophily in the Archaeon Sulfolobus solfataricus by Adaptive Evolution. Appl Environ Microbiol 82, 857–867

12. Payne, S., McCarthy, S., Johnson, T., North, E., and Blum, P. (2018) Nonmutational mechanism of inheritance in the Archaeon Sulfolobus solfataricus. Proc Natl Acad Sci U S A 115, 12271–12276

13. Dvorak, K., Payne, C. M., Chavarria, M., Ramsey, L., Dvorakova, B., Bernstein, H., Holubec, H., Sampliner, R. E., Guy, N., Condon, A., Bernstein, C., Green, S. B., Prasad, A., and Garewal, H. S. (2007) Bile acids in combination with low pH induce oxidative stress and oxidative DNA damage: relevance to the pathogenesis of Barrett’s oesophagus. Gut 56, 763–771

14. Selivanov, V. A., Zeak, J. A., Roca, J., Cascante, M., Trucco, M., and Votyakova, T. V. (2008) The role of external and matrix pH in mitochondrial reactive oxygen species generation. J Biol Chem 283, 29292–29300

15. Bird, A. P. (1986) CpG-rich islands and the function of DNA methylation. Nature 321, 209–213

16. Choli, T., Henning, P., Wittmann-Liebold, B., and Reinhardt, R. (1988) Isolation, characterization and microsequence analysis of a small basic methylated DNA-binding protein from the Archaebacterium, Sulfolobus solfataricus. Biochim Biophys Acta 950, 193–203

17. Guo, L., Feng, Y., Zhang, Z., Yao, H., Luo, Y., Wang, J., and Huang, L. (2008) Biochemical and structural characterization of Cren7, a novel chromatin protein conserved among Crenarchaea. Nucleic Acids Res 36, 1129–1137

18. Kreuz, S., and Fischle, W. (2016) Oxidative stress signaling to chromatin in health and disease. Epigenomics 8, 843–862

19. Niu, Y., DesMarais, T. L., Tong, Z., Yao, Y., and Costa, M. (2015) Oxidative stress alters global histone modification and DNA methylation. Free Radic Biol Med 82, 22–28

20. Hondorp, E. R., and Matthews, R. G. (2004) Oxidative stress inactivates cobalamin-independent methionine synthase (MetE) in Escherichia coli. PLoS Biol 2, e336

21. Banerjee, R. V., Frasca, V., Ballou, D. P., and Matthews, R. G. (1990) Participation of cob(I) alamin in the reaction catalyzed by methionine synthase from Escherichia coli: a steady-state and rapid reaction kinetic analysis. Biochemistry 29, 11101–11109

22. Mentch, S. J., Mehrmohamadi, M., Huang, L., Liu, X. J., Gupta, D., Mattocks, D., Padilla, P. G., Ables, G., Bamman, M. M., Thalacker-Mercer, A. E., Nichenametla, S. N., and Locasale, J. W. (2015) Histone Methylation Dynamics and Gene Regulation Occur through the Sensing of One-Carbon Metabolism. Cell Metab 22, 861–873

23. Martin, C., and Zhang, Y. (2005) The diverse functions of histone lysine methylation. Nat Rev Mol Cell Bio 6, 838–849

24. Kera, Y., Katoh, Y., Ohta, M., Matsumoto, M., Takano-Yamamoto, T., and Igarashi, K. (2013) Methionine Adenosyltransferase II-dependent Histone H3K9 Methylation at the COX-2 Gene Locus. Journal of Biological Chemistry 288, 13592–13601

25. Li, S., Swanson, S. K., Gogol, M., Florens, L., Washburn, M. P., Workman, J. L., and Suganuma, T. (2015) Serine and SAM Responsive Complex SESAME Regulates Histone Modification Crosstalk by Sensing Cellular Metabolism. Mol Cell 60, 408–421

26. Ye, C. Q., Sutter, B. M., Wang, Y., Kuang, Z., Zhao, X. Z., Yu, Y. H., and Tu, B. P. (2019) Demethylation of the Protein Phosphatase PP2A Promotes Demethylation of Histones to Enable Their Function as a Methyl Group Sink. Mol Cell 73, 1115–+

27. Vorontsov, E. A., Rensen, E., Prangishvili, D., Krupovic, M., and Chamot-Rooke, J. (2016) Abundant Lysine Methylation and N-Terminal Acetylation in Sulfolobus islandicus Revealed by Bottom-Up and Top-Down Proteomics. Mol Cell Proteomics 15, 3388–3404

28. Porcelli, M., Cacciapuoti, G., Carteni-Farina, M., and Gambacorta, A. (1988) S-adenosylmethionine synthetase in the thermophilic archaebacterium Sulfolobus solfataricus. Purification and characterization of two isoforms. Eur J Biochem 177, 273–280

29. Schelert, J., Dixit, V., Hoang, V., Simbahan, J., Drozda, M., and Blum, P. (2004) Occurrence and characterization of mercury resistance in the hyperthermophilic archaeon Sulfolobus solfataricus by use of gene disruption. J Bacteriol 186, 427–437

30. Rudrappa, D., Yao, A. I., White, D., Pavlik, B. J., Singh, R., Facciotti, M. T., and Blum, P. (2015) Identification of an archaeal mercury regulon by chromatin immunoprecipitation. Microbiology 161, 2423–2433

31. Struys, E. A., Jansen, E. E., de Meer, K., and Jakobs, C. (2000) Determination of *S*-adenosylmethionine and *S*-adenosylhomocysteine in plasma and cerebrospinal fluid by stable-isotope dilution tandem mass spectrometry. Clin Chem 46, 1650–1656

32. Lundback, T., Hansson, H., Knapp, S., Ladenstein, R., and Hard, T. (1998) Thermodynamic characterization of non-sequence-specific DNA-binding by the Sso7d protein from Sulfolobus solfataricus. J Mol Biol 276, 775–786

33. Kalichuk, V., Behar, G., Renodon-Corniere, A., Danovski, G., Obal, G., Barbet, J., Mouratou, B., and Pecorari, F. (2016) The archaeal “7 kDa DNA-binding” proteins: extended characterization of an old gifted family. Sci Rep 6, 37274

34. Zhang, Z., Zhao, M., Chen, Y., Wang, L., Liu, Q., Dong, Y., Gong, Y., and Huang, L. (2019) Architectural roles of Cren7 in folding crenarchaeal chromatin filament. Mol Microbiol 111, 556–569

35. Martinez, A., and Kolter, R. (1997) Protection of DNA during oxidative stress by the nonspecific DNA-binding protein Dps. J Bacteriol 179, 5188–5194

36. Nguyen-Duc, T., van Oeffelen, L., Song, N., Hassanzadeh-Ghassabeh, G., Muyldermans, S., Charlier, D., and Peeters, E. (2013) The genome-wide binding profile of the Sulfolobus solfataricus transcription factor Ss-LrpB shows binding events beyond direct transcription regulation. BMC Genomics 14, 828

37. Nguyen, A. T., and Zhang, Y. (2011) The diverse functions of Dot1 and H3K79 methylation. Genes Dev 25, 1345–1358

38. Huang, B. X., and Kim, H. Y. (2013) Effective identification of Akt interacting proteins by two-step chemical crosslinking, co-immunoprecipitation and mass spectrometry. PLoS One 8, e61430

39. Ishioka, K., Fukuoh, A., Iwasaki, H., Nakata, A., and Shinagawa, H. (1998) Abortive recombination in Escherichia coli ruv mutants blocks chromosome partitioning. Genes Cells 3, 209–220

40. Bergerat, A., de Massy, B., Gadelle, D., Varoutas, P. C., Nicolas, A., and Forterre, P. (1997) An atypical topoisomerase II from Archaea with implications for meiotic recombination. Nature 386, 414–417

41. Jolma, A., Yin, Y. M., Nitta, K. R., Dave, K., Popov, A., Taipale, M., Enge, M., Kivioja, T., Morgunova, E., and Taipale, J. (2015) DNA-dependent formation of transcription factor pairs alters their binding specificity. Nature 527, 384–+

42. Funnell, A. P., and Crossley, M. (2012) Homo- and heterodimerization in transcriptional regulation. Adv Exp Med Biol 747, 105–121

43. Kohen, R., and Nyska, A. (2002) Oxidation of biological systems: oxidative stress phenomena, antioxidants, redox reactions, and methods for their quantification. Toxicol Pathol 30, 620–650

44. Salvatore, F., Zappia, V., and Shapiro, S. K. (1968) Quantitative analysis of S-adenosylhomocysteine in liver. Biochim Biophys Acta 158, 461–464

45. Lu, Z. J., and Markham, G. D. (2002) Enzymatic properties of S-adenosylmethionine synthetase from the archaeon Methanococcus jannaschii. J Biol Chem 277, 16624–16631

46. Sweetlove, L. J., and Fernie, A. R. (2018) The role of dynamic enzyme assemblies and substrate channelling in metabolic regulation. Nat Commun 9, 2136

47. Zhang, Z., Gong, Y., Guo, L., Jiang, T., and Huang, L. (2010) Structural insights into the interaction of the crenarchaeal chromatin protein Cren7 with DNA. Mol Microbiol 76, 749–759

48. Chu, Y., Zhang, Z., Wang, Q., Luo, Y., and Huang, L. (2012) Identification and characterization of a highly conserved crenarchaeal protein lysine methyltransferase with broad substrate specificity. J Bacteriol 194, 6917–6926

49. Boukouris, A. E., Zervopoulos, S. D., and Michelakis, E. D. (2016) Metabolic Enzymes Moonlighting in the Nucleus: Metabolic Regulation of Gene Transcription. Trends Biochem Sci 41, 712–730

50. Ekman, D., Bjorklund, A. K., Frey-Skott, J., and Elofsson, A. (2005) Multi-domain proteins in the three kingdoms of life: orphan domains and other unassigned regions. J Mol Biol 348, 231–243

51. Galperin, M. Y., Makarova, K. S., Wolf, Y. I., and Koonin, E. V. (2018) Phyletic Distribution and Lineage-Specific Domain Architectures of Archaeal Two-Component Signal Transduction Systems. J Bacteriol 200

52. Melnikov, S., Kwok, H. S., Manakongtreecheep, K., van den Elzen, A., Thoreen, C. C., and Soll, D. (2019) Archaeal ribosomal proteins possess nuclear localization signal-type motifs: implications for the origin of the cell nucleus. Mol Biol Evol

53. Lyons, T. W., Reinhard, C. T., and Planavsky, N. J. (2014) The rise of oxygen in Earth’s early ocean and atmosphere. Nature 506, 307–315

