## Supplementary material for "Conservation of an Ancient Oxidation Response That Controls Archaeal Epigenetic Traits Through Chromatin Protein Networks": SI Appendix

### Supplementary Methods

**Archaeal strains and cultivation**. *S. solfataricus* was cultured at 80°C in screw-cap glass flasks with shaking as described previously (1-3) in Allen's basal salts (1) as modified (4), supplemented with 0.2% (w/v) tryptone. PBL2025 was cultivated at pH 3.0 while SARC-I was cultivated at both pH 3.0 and pH 1.0. For biomass used for chromatin immunoprecipitation (ChIP), samples were prepared from 500mL cultures.

**Immunoprecipitation for ChIP-Seq**. Crosslinked cell pellets were resuspended in 200uL of lysis buffer (50mM HEPES-KOH, 140mM NaCl, 1mM EDTA, 1% Triton X-100, 0.1% Na-deoxycholate, pH 7.5) and sonicated using a FB705 cup-horn sonicator (Thermofisher, Walthan, MA). The cell lysates were clarified for 20 min at 16,000 x g prior to their use in ChIP. Cell lysate was combined with 20µL protein A-conjugated Sepharose 4B beads (5) (Life technologies cat# 101041) that had been pre-incubated with 20µL Cren7 polyclonal antibody serum or 100µl Sso7d antibody serum for 2hr at room temperature with agitation and washed twice with 1mL PBS. The bead-lysate mixture was incubated with agitation for 2hr at room temperature, and then washed three times with 1mL of PBS. ChIP samples were eluted and RNAse treated as described (6). DNA samples were then prepared to make libraries using phenol chloroform extraction followed by ethanol precipitation. Background controls of whole cell extract (WCE) were included for each sample and were later used as ChIP-seq controls (6).

**ChIP-seq.** Barcode sequences are provided in SI Appendix, Table S5. Samples were used to template PCR amplification for 12 cycles, and PCR products were visualized and quantified using a bioanalyzer (Agilent, Santa Clara, CA) and a high-sensitivity DNA ChIP (Agilent cat. # 5067-4626). ChIP and background libraries were pooled in equimolar concentrations and submitted for Illumina HiSeq3000 SR50 sequencing, yielding about 300 million 50bp reads.

**ChIP-Seq data analysis.** Quality controlled libraries were aligned to the SULG genome (NZ_CP033235.1) using Bowtie (7) and converted into sorted BAM files using the SAMtools package (8). Alignment quality for each library are described in SI Appendix, Table S6. The DeepTools package (9) multiBAMSummary, plotCorrelation and computeGCBias tools were used to calculate the reproducibility of duplicate ChIP samples (Pearsons correlation coefficient of >0.99) and to determine library quality. Normalized ChIP-seq coverage files were generated using DeepTools bamCompare, which computed the difference in the number of reads for the WCE background control libraries and their respective ChIP samples while normalizing for sequencing depth (10,11). Resulting Bigwig data was visualized using IGV (12,13) and high affinity peaks were recorded. The threshold for identifying high affinity peaks was calculated as the average reads per bp across the genome multiplied by 100. The annotation of bound ORFs and the relative location of peaks within the ORF were recorded. MEME-ChIP (v5.0.5) was used to search for conserved binding motifs across a 100 bp region located at the center of sequences associated with ChIPseq peaks, using standard settings (14). Regions of GC bias were searched for using methods described previously (15).

**Immunoprecipitation for AP-MS.** Resuspended cell pellets were sonicated to about a 1000bp DNA fragment size. Cell lysates were combined with 50µL of Protein A conjugated Sepharose 4B beads incubated with 50µL of antibody serum. After incubation, lysate-bead slurries were washed 2x with 1mL PBS. To remove DNA that might link distal protein complexes, beads were treated with 5µL DNAse (Thermofisher, cat# EN0521). Samples were incubated for 30 min at 37°C, washed 2x with 1mL PBS and then re-digested with 2µL DNAse to ensure complete removal of exposed DNA. Samples were washed 3x with 1mL PBS and protein was eluted in 200µL 50mM Tris-HCl pH 8, 10mM EDTA, 5% SDS and then incubated overnight at 67°C to reverse crosslinking. Complete DNA digestion was validated using DNA gel electrophoresis and Qubit 2 (Thermofisher, Walthan, MA) measurement compared to undigested samples. To control for background precipitations, control samples immunoprecipitated without antibody serum and with pre-immune serum were included (16).

**LC-MS/MS of ChIP samples.** Protein samples were digested with Trypsin Gold (Promega) and purified using an S-Trap mini column (Protifi) following the manufacturer's protocol. Samples were dried using a Speed Vac, stored at -80ºC and then reconstituted in 50 µL 5% formic acid in water prior to LC injection. Digested samples were spiked with Yeast Alcohol Dehydrogenase digest (Waters, Cat# 186002325) as a quantification reference and subjected to nanoLC-MS/MS analysis on a Sciex 6600 Triple-TOF with DuoSpray source and Calibrant Delivery System (CDS) coupled to an Ultimate 3000 nano-LC system (Dionex Corporation, USA). The peptides were separated on a C18 Pep Map column (1 mm ID x 150 mm L, 3 µm particle size, 100Å pore, Dionex) by applying an acetonitrile (ACN) gradient (ACN plus 0.1% formic acid, 60 min gradient of 0-40%, then 2 min gradient of 30-40%, then held at 40% for 5 min) at a flow rate of 50 µL/min, and introduced into the mass spectrometer using the nanospray source. All MS methods for the 6600 Triple-TOF used the information dependent acquisition mode. The 6600 Triple-TOF was operated with the following parameters: nano spray voltage, positive mode; survey scan range, 400-1250 m/z; MS/MS scan range, 100-1500 m/z; 48 MS/MS scans/cycle; rolling collision energy.

**Quantitative AP-MS data analysis.** The MS/MS spectra were searched against the *Sulfolobus solfataricus* proteome from UniProt using ProteinMetrics Byonic and Byologic (v3.2). Database search criteria were as follows: enzyme: Trypsin, missed cleavages: 2; mass: monoisotopic; fixed modification: carbamidomethyl (C); variable modification: acetylation (K) and methylation (K); precursor tolerance: 20 ppm; product ion tolerance: 60 ppm. The Byologic Top3 algorithm was used to quantify protein abundance using the spiked ADH digest as a reference. Protein abundance was normalized between replicates to the abundance of the target chromatin protein. Protein abundance was calculated as the net increase in experimental sample yield compared to a no-antibody background control. Proteins whose abundance was 5-fold greater in the experimental samples compared to Day 0 antibody background controls were considered to be enriched by ChIP and associated with Cren7 or Sso7D. Proteins whose abundance varied > 50% between duplicates were filtered out for having experimental noise. To avoid false negatives resulting from free peptides, proteins required at least 25% coverage in one sample. For differential abundance analysis, changes resulting from expression differences and not protein affinity changes were removed. Protein identity and function were refined by using NCBI's BLAST software (17), the Conserved Domain Database (18), and KEGG pathways (19).

**SAM extraction.** SAM was extracted by solid phase extraction (SPE) as in Struys, et al. (20), with modifications as follows. 15OD's of cells from mid-exponential phase cultures were deproteinized with 2 mL 10% perchloric acid per 100 mg (wet weight) cells and incubation on ice for 1 hour. The internal standard, methylated adenosine (Abcam, cat# ab145715), was added to each sample at a concentration of 10 µM (21). Samples were diluted with 1 mL of water and neutralized with 0.5 mL of 1 M sodium phosphate pH 11.5.The Waters Oasis WCX SPE cartridges were conditioned with 1 mL of methanol, followed by 2 mL water, 750 µL of 100 mM sodium phosphate pH 7.0, and 1 mL of water. After sample application, cartridges were washed with 2 mL water. SAM was eluted with two 600 µL volumes of SPE elution solvent (50:49:1 methanol: water: trifluoracetic acid). The eluted samples evaporated to dryness in a SpeedVac SC110 (Savant Instruments, Holbrook, NY) concentrator at room temperature. The samples were reconstituted twice with 100 µL of mobile phase solvent A and all 200µL was transferred to glass HPLC vials with limited volume inserts. The HPLC fractionation of SAM was adapted from Stabler et al 2004 (22), with modifications. A 100 mm Thermo Aquasil C18 column (2.1 mm inner diameter and 3 µm particles) was maintained at 30°C using an Agilent 1200 Series Rapid Resolution HPLC (Agilent Technologies, Santa Clara, CA) equipped with a binary pump. 5 µL was injected for each sample, and the autosampler needle was washed in 85% water, 15% methanol between injections. A diode array detector measured SAM absorbance at 260nm.

**Data availability**. The ChIPseq raw and processed data files are available at the Gene Expression Omnibus. For Cren7 data, the accession numbers are: SULG at pH 3 (XXX, XXX, XXX), SARC-I at pH 3 (XXX, XXX, XXX) and SARC-I at pH 3 (XXX, XXX, XXX). For Sso7D data, the accession numbers are: SULG at pH 3 (XXX, XXX, XXX), SARC-I at pH 3 (XXX, XXX, XXX) and SARC-I at pH 3 (XXX, XXX, XXX).

**Supplemental figures**

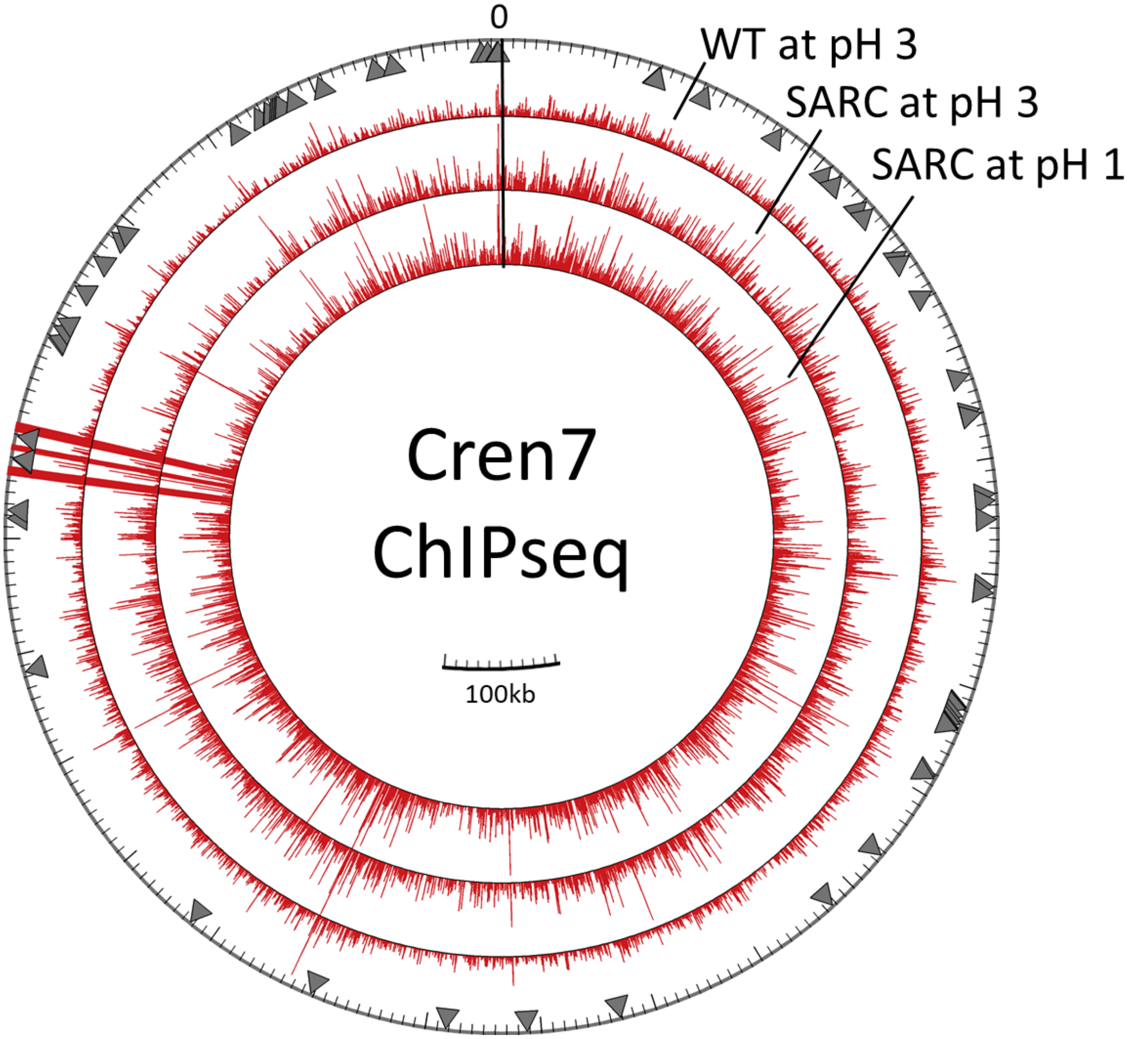

**Fig. S1. Distribution of Cren7 chromatin proteins in the *S. solfataricus* genome.** Bar height indicates ChIP-seq read depth averaged across a 50bp window after subtracting background DNA. The maximum displayed peak height is 3000 reads. Reads are mapped to the SULG genome. Grey triangles indicate the location of SARC transcriptome genes.

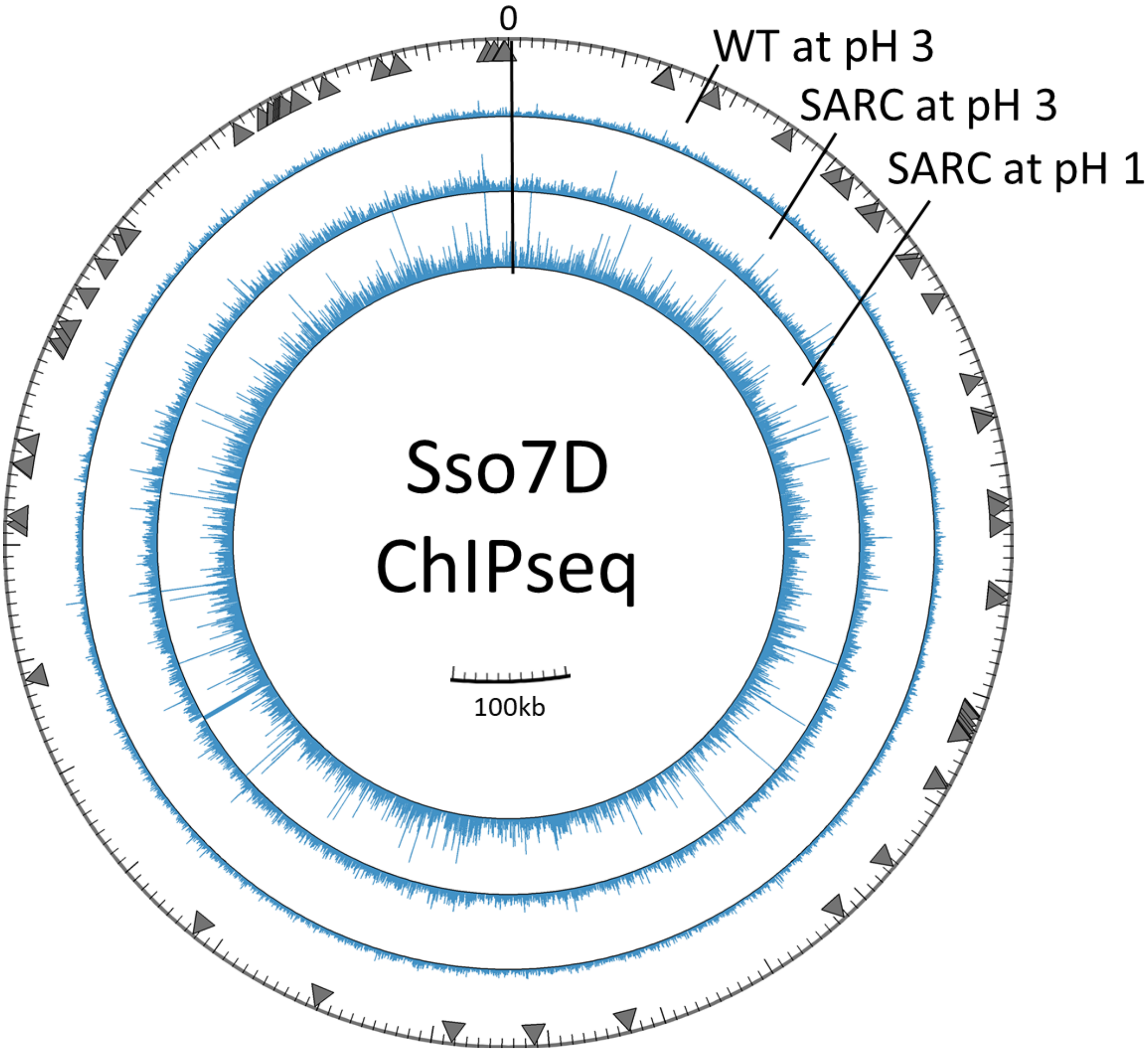

**Fig. S2. Distribution of Sso7D chromatin proteins in the *S. solfataricus* genome.** Bar height indicates ChIP-seq read depth averaged across a 50bp window after subtracting background DNA. The maximum displayed peak height is 1000 reads. Reads are mapped to the SULG genome. Grey triangles indicate the location of SARC transcriptome genes.

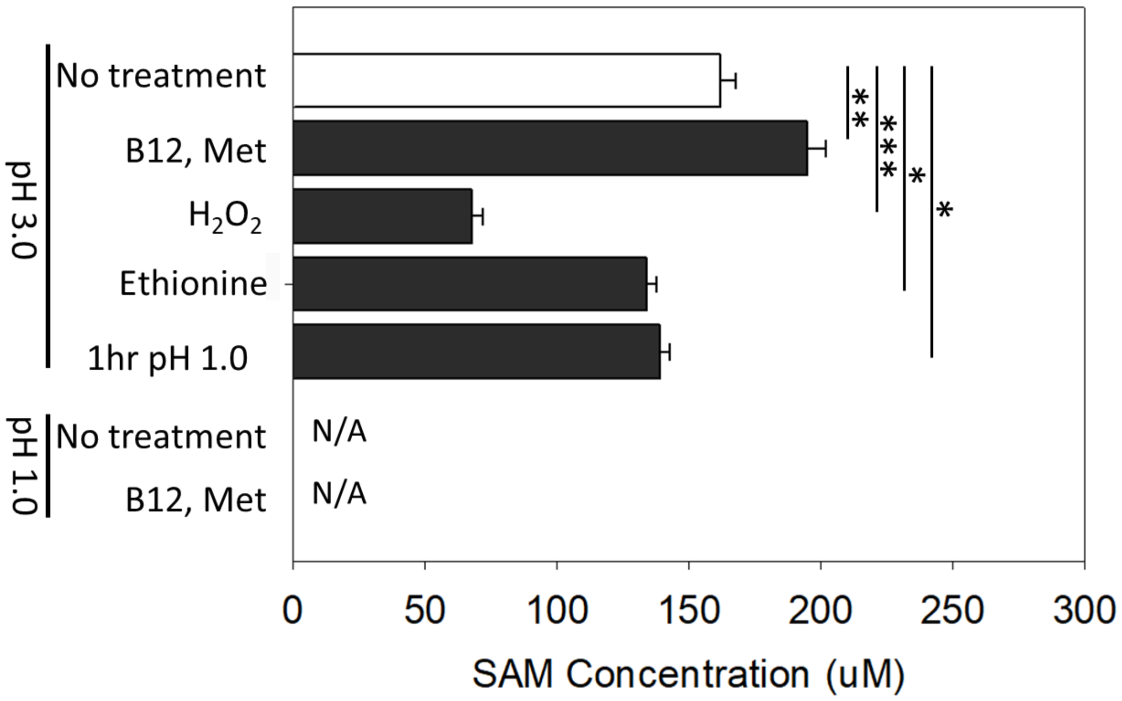

**Fig. S3. Effect of treatments on S-adenosylmethionine (SAM) abundance in Wild Type.** SULG was passaged 3x at a reduction potential of pH 3.0 (504mV). pH 3.0 cultures were treated with oxidative stress (120uM H2O2) for 1hr prior to SAM extraction, with 1mM ethionine during culture, with 500nM Vitamin B12 and 10mM methionine for 12hrs prior to SAM extraction, or transferred to pH 1.0 media for 1hr prior to extraction. N/A indicates that treatments that were not possible for the SULG strain. n = 4 for all conditions. (*) $P < 0.05$, (**) $P < 0.01$, (***) $P < 0.001$, Students T-test

**Supplementary Tables**

**Table S1. High affinity peaks in the Cren7 ChIPseq dataset.**

| SULA ORF | Protein Accession # | Annotation | Peak location within ORF (Start = 1-30%, mid = 31-69%, end = 70-100%) |
|---|---|---|---|
| SULA_0024 | A0A0E3K9V8 | Uncharacterized protein | Start-Mid |
| SULA_0053 | N/A | Hypothetical protein | Start-Mid |
| SULA_0055 | A0A0E3MEX0 | Uncharacterized protein | End |
| SULA_0058 | A0A0E3K9X1 | Nucleoside hydrolase | Start-Mid |
| SULA_0093 | A0A0E3MFV9 | Cysteine--tRNA ligase | Mid |
| SULA_0101 | A0A0E3MBP1 | MFS transporter | Intergenic |
| SULA_0102 | N/A | SDR family oxidoreductase | |
| SULA_0103 | A0A0E3JVP9 | Exodeoxyribonuclease III | Start |
| SULA_0108 | A0A0E3GVU9 | Cobalt-precorrin-6Y C(5)-methyltransferase (CbiE) | End |
| SULA_0110 | A0A0E3MBP3 | Precorrin-4 C(11)-methyltransferase (CbiL) | Intergenic |
| SULA_0111 | A0A0E3GSS1 | cobalt-precorrin-2 C(20)-methyltransferase (CbiF) | |
| SULA_0142 | A0A0E3M9F1 | Integrase | Intergenic |
| SULA_0155 | A0A0E3JZP3 | Phosphoesterase | Start |
| SULA_0159 | A0A0E3JSQ8 | DUF2070 family protein | Mid |
| SULA_0180 | A0A0E3K6B5 | Acetyl-CoA C-acyltransferase | Mid |
| SULA_0205 | A0A0E3K7J9 | Hydrolase | Mid |
| SULA_0254 | A0A0E3KB42 | Inorganic phosphate transporter (Phosphate permease) | Start |
| SULA_0265 | A0A0E3MHK4 | Deoxyribodipyrimidine photo-lyase | End |
| SULA_0300 | A0A0E3KB53 | Acyl-CoA dehydrogenase | Mid |
| SULA_0305 | A0A0E3K4W1 | 4-hydroxybutyryl-CoA dehydratase | Mid |
| SULA_0325 | A0A0E3MHM9 | FAD-dependent oxidoreductase | Mid |
| SULA_0335 | A0A0E3K6I3 | Phosphoenolpyruvate carboxykinase [GTP] | Mid-End |
| SULA_0342 | N/A | IS1182 Transposase | End |
| SULA_0353 | A0A0E3GSW4 | Peptidase S53 | Mid |
| SULA_0403 | A0A0E3GUG2 | Protein-L-isoaspartate O-methyltransferase | Mid |
| SULA_0436 | A0A3G8DZ91 | DUF1641 domain-containing protein | Mid |
| SULA_0447 | A0A0E3MG97 | Glycerol-3-phosphate dehydrogenase | Mid |
| SULA_0474 | A0A0E3JWD4 | ABC transporter ATP-binding protein | Mid-End |
| SULA_0495 | A0A0E3GW17 | CBS domain-containing protein | End |
| SULA_0502 | A0A0E3GSZ0 | TenA family transcriptional regulator | Start |
| SULA_0515 | A0A3G8EHC5 | ABC transporter ATP-binding protein | End |
| SULA_0544 | A0A0E3JWH4 | Glycoside hydrolase family 15 protein | End |
| SULA_0568 | A0A0E3GUJ2 | 4Fe-4S dicluster domain-containing protein | Mid |
| SULA_0693 | A0A0E3KAL5 | Lycopene cyclase | Intergenic |
| SULA_0694 | A0A0E3MC61 | Phytoene synthase | |
| SULA_0728 | A0A0E3GW58 | 2-hydroxyhepta-2,4-diene-1,7-dioate isomerase | End |
| SULA_0731 | A0A0E3MBG5 | DUF3782 domain-containing protein | End |
| SULA_0739 | A0A0E3KAN3 | Uncharacterized protein | Start |
| SULA_0743 | A0A0E3K6Z3 | ABC transporter ATP-binding protein | Mid |
| SULA_0745 | A0A0E3K5C6 | ABC transporter permease | Intergenic |
| SULA_0746 | A0A0E3GT33 | ABC transporter substrate-binding protein | |
| SULA_0758 | A0A0E3KAP2 | Quinol oxidase subunit 1 (Quinol oxidase subunit 1/3) | Start |

| | | | |
|---|---|---|---|
| SULA_0758 | A0A0E3KAP2 | Quinol oxidase subunit 1 (Quinol oxidase subunit 1/3) | Mid |
| SULA_0834 | N/A | ABC transporter | Mid |
| SULA_0862 | A0A0E3MBJ5 | ABC transporter ATP-binding protein | Intergenic |
| SULA_0863 | A0A0E3MF58 | Peroxiredoxin | |
| SULA_0878 | A0A0E3K896 | Uncharacterized protein | End |
| SULA_0883 | A0A0E3KAV3 | Metal-dependent carboxypeptidase | Mid-End |
| SULA_0901 | A0A0E3MDW3 | HD domain-containing protein | End |
| SULA_0940 | A0A0E3K5H5 | Uncharacterized protein | Start |
| SULA_0943 | A0A0E3JTH1 | ABC transporter ATP-binding protein | Intergenic |
| SULA_0944 | A0A0E3K792 | tRNA (N6-threonylcarbamoyladenosine(37)-N6)-methyltransferase TrmO | |
| SULA_0971 | A0A0E3MFE3 | Oxidoreductase | Mid |
| SULA_0975 | A0A0E3K7A3 | CBS domain-containing protein | Intergenic |
| SULA_0976 | A0A0E3KB05 | Endoribonuclease (RidA family protein) | |
| SULA_0988 | A0A0E3JTI6 | Gamma-glutamyltransferase | Start |
| SULA_0997 | A0A0E3GU65 | CBS domain-containing protein | all |
| SULA_1083 | A0A0E3MA25 | MEMO1 family protein | Mid |
| SULA_1103 | A0A0E3K0B0 | PDZ domain-containing protein | End |
| SULA_1113 | A0A0E3MA41 | Methionine aminopeptidase | Start |
| SULA_1117 | P95959 | Alpha/beta hydrolase | Mid |
| SULA_1199 | A0A0E3MA51 | Porphobilinogen deaminase (hemC) | Intergenic |
| SULA_1200 | A0A0E3K8L7 | Uroporphyrinogen III methyltransferase (hemD) | |
| SULA_1205 | A0A0E3K5P8 | MBL fold metallo-hydrolase (RNA procession exonuclease-like protein) | Intergenic |
| SULA_1206 | A0A0E3MIT8 | DNA ligase | |
| SULA_1216 | A0A3G8DFX3 | S-adenosylmethionine synthase | End |
| SULA_1255 | A0A0E3MKK3 | Thiamine-monophosphate kinase | Mid |
| SULA_1257 | A0A0E3GUW0 | Adenylosuccinate lyase | Start |
| SULA_03245 | #N/A | Integrase (pseudogene) | Intergenic |
| SULA_1307 | A0A3G8EUD1 | AbrB/MazE/SpoVT family DNA-binding domain-containing protein | |
| SULA_1340 | A0A0E3KBE2 | Flavin-dependent thymidylate synthase | Start |
| SULA_1348 | A0A0E3MIY8 | Oxidoreductase | Mid |
| SULA_1365 | A0A0E3KC99 | Prefoldin subunit alpha (GimC subunit alpha) | Start |
| SULA_1412 | A0A0E3K770 | tRNA(Met) cytidine acetyltransferase TmcA | Mid |
| SULA_1478 | A0A0E3GWI8 | DUF72 domain-containing protein | Intergenic |
| SULA_1479 | A0A0E3JXJ6 | LysE family translocator | |
| SULA_1510 | A0A0E3MC21 | ABC transporter ATP-binding protein | End |
| SULA_1656 | A0A0E3MC63 | ATP synthase subunit F (V-type ATP synthase subunit F) | End |
| SULA_1671 | Q9UWY1 | Acetolactate synthase | Intergenic |
| SULA_1672 | A0A0E3MHM7 | Magnesium-dependent phosphatase-1 | |
| SULA_1684 | A0A0E3MHN1 | Histidinol-phosphate aminotransferase | Mid |
| SULA_1693 | O33779 | Metal-sulfur cluster assembly factor | Start |
| SULA_1705 | A0A0E3GV39 | Aspartate carbamoyltransferase regulatory chain | End |
| SULA_1717 | A0A0E3GTK2 | Methionine--tRNA ligase subunit beta | Intergenic |
| SULA_1718 | A0A0E3JU83 | Phosphoribosylaminoimidazole-succinocarboxamide synthase | |
| SULA_1720 | A0A0E3MC78 | Phosphoribosylformylglycinamidine synthase subunit PurQ | Intergenic |
| SULA_1721 | A0A0E3GWN1 | Phosphoribosylformylglycinamidine synthase subunit PurL | |
| SULA_1723 | A0A0E3JVD7 | Amidophosphoribosyltransferase | Start |

| | | | |
|---|---|---|---|
| SULA_1725 | A0A0E3MG02 | Phosphoribosylformylglycinamidine cyclo-ligase | Start |
| SULA_1727 | A0A0E3K8A1 | CopG family transcriptional regulator | Intergenic |
| SULA_1728 | A0A0E3GWN2 | Argininosuccinate synthase | |
| SULA_1729 | A0A0E3GTK4 | Argininosuccinate lyase | Start |
| SULA_1743 | Q9UX47 | Homoserine dehydrogenase | Start |
| SULA_1758 | Q9UX63 | CBS domain-containing protein | Intergenic |
| SULA_1759 | Q9UX64 | Molybdopterin-guanine dinucleotide biosynthesis protein B | |
| SULA_1763 | Q9UX68 | Amidohydrolase | Mid |
| SULA_1769 | Q9UX75 | ABC transporter permease | Mid |
| SULA_1806 | A0A0E3ME04 | Isoleucine--tRNA ligase | Start |
| SULA_1841 | A0A0E3JXV6 | Polyamine aminopropyltransferase | Start |
| SULA_1875 | A0A0E3JUB5 | IS640 transposase | Intergenic |
| SULA_1894 | A0A0E3GTN3 | Threonine synthase | Mid |
| SULA_1895 | A0A0E3MFP2 | UbiD family decarboxylase | End |
| SULA_1905 | A0A0E3JUC1 | Anthranilate synthase component 1 | Mid |
| SULA_1906 | A0A0E3K8G6 | Aminodeoxychorismate/anthranilate synthase component II | Intergenic |
| SULA_1907 | A0A0E3JXX5 | Indole-3-glycerol phosphate synthase | |
| SULA_1908 | A0A0E3KC25 | Aminotransferase | Intergenic |
| SULA_1909 | A0A0E3GV75 | Valine--tRNA ligase | |
| SULA_1984 | A0A0E3MDS4 | 3-hydroxyisobutyrate dehydrogenase | Mid |
| SULA_1999 | A0A0E3MJN4 | NDP-sugar synthase | Intergenic |
| SULA_2000 | A0A0E3K136 | Glucan 1,3-alpha-glucosidase | |
| SULA_2000 | A0A0E3K136 | Glucan 1,3-alpha-glucosidase | End |
| SULA_2030 | A0A0E3MJL1 | Uncharacterized protein | Intergenic |
| SULA_2031 | A0A0E3GWT5 | TatD family deoxyribonuclease | |
| SULA_2034 | A0A0E3MAU5 | Extracellular solute-binding protein | End |
| SULA_2052 | A0A0E3JY42 | DNA primase small subunit PriS | Intergenic |
| SULA_2053 | A0A0E3KC79 | Uncharacterized protein | |
| SULA_2082 | A0A0E3MI23 | DSBA oxidoreductase | Mid |
| SULA_2091 | A0A0E3MEW0 | CoA transferase | Intergenic |
| SULA_2092 | A0A0E3KD43 | Uncharacterized protein | |
| SULA_2094 | A0A0E3KC94 | VWA domain-containing protein | Start |
| SULA_2119 | A0A0E3KD47 | Biotin synthase (Radical SAM protein) | Intergenic |
| SULA_2120 | A0A0E3GVB1 | Uncharacterized protein | |
| SULA_2136 | A0A0E3GTS5 | Uncharacterized protein | End |
| SULA_2150 | A0A0E3JZH9 | Uncharacterized protein | End |
| SULA_2153 | A0A0E3GTS8 | Acyl-CoA dehydrogenase | End |
| SULA_2169 | A0A0E3KCF7 | Glycoside hydrolase | Start |
| SULA_2170 | A0A0E3GVC0 | APC family permease | Mid |
| SULA_2174 | A0A0E3K692 | Aminotransferase | Mid |
| SULA_2233 | A4ZY71 | Multicomponent monooxygenase system protein E | Mid |
| SULA_2252 | A0A0E3JVF8 | LOG family protein | End |
| SULA_2254 | A0A0E3KCI7 | Uncharacterized protein | Mid |
| SULA_2258 | A0A0E3K6C4 | RNA helicase | End |
| SULA_2275 | A0A0E3KCJ6 | ABC transporter substrate-binding protein | Mid |
| SULA_2277 | A0A0E3K9X3 | Glycosyltransferase | End |
| SULA_2310 | A0A0E3GVE3 | Type I-A CRISPR-associated protein Cas7/Csa2 | Mid |
| SULA_2313 | A0A0E3GWY2 | CRISPR-associated helicase Cas3 | Mid |

| | | | |
|---|---|---|---|
| SULA_2320 | A0A0E3MCN8 | Acylaminoacyl peptidase (S9 family peptidase) | Mid |
| SULA_0292 | A0A0E3GSW3 | IS5-like element ISC1234 family transposase | Intergenic |
| SULA_2322 | A0A0E3MDF7 | CRISPR-associated exonuclease Cas4 | |
| SULA_2434 | A0A0E3GX01 | 4Fe-4S ferredoxin | Start |
| SULA_2434 | A0A0E3GX01 | 4Fe-4S ferredoxin | Mid |
| SULA_2436 | A0A0E3MJY7 | Uncharacterized protein | Start |
| SULA_2500 | A0A0E3MEG0 | Amine oxidase | Mid |
| SULA_2625 | A0A0E3GU03 | Uncharacterized protein | Intergenic |
| SULA_2626 | A0A0E3KDL3 | FtsX-like permease family protein | |
| SULA_2631 | A0A0E3K6R8 | Uncharacterized protein | Start |
| SULA_2710 | A0A0E3GVK5 | ATP-binding cassette domain-containing protein (Cobalt ABC transporter ATP-binding protein) | End |
| SULA_0352 | A0A0E3MIM1 | DNA-binding protein Sso7d | Intergenic |
| SULA_2728 | A0A0E3GX46 | MFS transporter | End |
| SULA_2821 | P95860 | Orf c06013 protein (Phosphoesterase) | End |
| SULA_2828 | P95868 | Glycogen debranching enzyme GlgX | Mid |
| SULA_2828 | P95868 | Glycogen debranching enzyme GlgX | End |
| SULA_2868 | P95901 | MFS transporter | Mid |
| SULA_2871 | P95904 | Pyruvate-ferredoxin oxidoreductase beta-2 | Mid |
| SULA_2921 | A0A0E3MHT7 | Aspartate-semialdehyde dehydrogenase | Mid |
| SULA_2923 | A0A0E3MKH6 | Uncharacterized protein | Intergenic |
| SULA_2924 | A0A3G8DRM1 | Peptidase S53 | |

**Table S2. High affinity peaks in the Sso7D ChIPseq dataset.**

| SULA ORF | Protein Accession # | Annotation | Peak location within ORF (Start = 1-30%, mid = 31-69%, end = 70-100%) |
|---|---|---|---|
| SULA_0028 | A0A0E3MAU4 | Radical SAM/SPASM domain-containing protein | Mid |
| SULA_0138 | A0A0E3MC65 | ISC1439B family transposase | Start |
| SULA_0149 | A0A0E3KA03 | Geranylgeranyl pyrophosphate synthase | Start |
| SULA_0158 | A0A0E3KB01 | Geranylgeranyl hydrogenase | Mid |
| SULA_0172 | A0A0E3KB06 | Aminotransferase class I/II-fold pyridoxal phosphate-dependent enzyme | Start |
| SULA_0221 | A0A0E3MG10 | Methylmalonyl-CoA mutase | Start-Mid |
| SULA_0258 | A0A0E3MEI0 | Acyl-CoA carboxylase subunit beta | Mid |
| SULA_0325 | A0A0E3MHM9 | FAD-dependent oxidoreductase | Start |
| SULA_0325 | A0A0E3KB57 | MFS transporter | Start |
| SULA_0344 | A0A0E3MC65 | ISC1439 transposase | Start-Mid |
| SULA_0453 | A0A3G8DE51 | YHS domain-containing protein | Intergenic |
| SULA_0454 | A0A0E3K517 | AsnC family transcriptional regulator | |
| SULA_0521 | A0A0E3KAD4 | Aminotransferase | Mid |
| SULA_0550 | A0A0E3JWH5 | Uncharacterized protein | End |
| SULA_0551 | A0A0E3K011 | Bacteriocin | |
| SULA_0561 | A0A0E3KBF1 | Pyruvate synthase | Mid |
| SULA_0602 | A0A0E3GW36 | Twin-arginine translocation signal domain-containing protein | End |
| SULA_0830 | A0A0E3MIU5 | IS1-like element ISC1173a family transposase | End |
| SULA_0908 | A0A0E3GT61 | MBL fold metallo-hydrolase | Start-Mid |
| SULA_0925 | A0A0E3GT64 | DUF981 family protein | Mid |
| SULA_0989 | N/A | Transposase (pseudogene) | Intergenic |
| SULA_0990 | N/A | Hypothetical protein | |
| SULA_1060 | A0A0E3JVB2 | Oxidase | Mid |
| SULA_1157 | A0A0E3MH25 | SSV1-like integrase, C-terminal domain) | Start |
| SULA_1277 | N/A | RNAse RNA A subunit | Mid |
| SULA_1576 | A0A0E3MCU1 | DEAD/DEAH box helicase | Start |
| SULA_1590 | A0A0E3MHJ6 | ATP-binding protein | Mid |
| SULA_03325 | A0A3G8DG08 | Ribbon-helix-helix protein, CopG family | Mid |
| SULA_1652 | Q9UWW0 | CDP-alcohol phosphatidyltransferase | Intergenic |
| SULA_1653 | A0A0E3MCY1 | DNA primase large subunit PriL | |
| SULA_1671 | Q9UWY1 | Acetolactate synthase | Start |
| SULA_1784 | A0A0E3MDL9 | 30S ribosomal protein S5 | Mid |
| SULA_1801 | A0A0E3MG43 | 50S ribosomal protein L2 | Mid |
| SULA_1923 | A0A0E3K644 | Cell division protein CdvA | Start |
| SULA_1929 | A0A0E3KC33 | Glycine cleavage system P-protein subunit 2 | Mid |
| SULA_1937 | A0A0E3MG73 | Fe-S cluster assembly protein SufB | Mid |
| SULA_2019 | N/A | rRNA 16S | All |
| SULA_2020 | N/A | rRNA 23S | All |
| SULA_2048 | A0A0E3MEV0 | RNA methyltransferase | Mid |
| SULA_2090 | A0A0E3K8N9 | Glutaconate CoA-transferase, subunit A (GctA) | End |
| SULA_2095 | A0A0E3MGU6 | FHA domain-containing protein | Start-Mid |
| SULA_2096 | A0A0E3KAT7 | Serine/threonine phosphatase | |

| | | | |
|---|---|---|---|
| SULA_2119 | A0A0E3KD47 | Biotin synthase (Radical SAM protein) | Start |
| SULA_2139 | A0A0E3KCE5 | MFS transporter | Mid |
| SULA_2140 | A0A0E3JUH7 | Peptidase S53 | Start-Mid |
| SULA_03475 | N/A | Pseudogene | Mid |
| SULA_2187 | A0A0E3MDC6 | FAD-dependent oxidoreductase | End |
| SULA_2304 | A0A0E3MH28 | CRISPR-associated exonuclease Cas4 | Mid |
| SULA_2328 | A0A0E3K1C8 | Glutamate dehydrogenase | Start |
| SULA_2392 | A0A0E3MGH4 | ATP-NAD kinase | Start |
| SULA_2405 | N/A | Pseudogene | Mid |
| SULA_2484 | A0A0E3GVH2 | ISH3-like element ISC1439B family transposase | Mid |
| SULA_2489 | A0A0E3MDQ2 | ISH3-like element ISC1439A family transposase | Mid |
| SULA_2576 | A0A0E3JVH5 | DoxX family membrane protein | Mid |
| SULG_13155 | A0A3G8EDC1 | Hypothetical protein | Intergenic |
| SULA_2587 | A0A0E3GVI7 | Hypothetical protein | |
| SULA_2653 | A0A0E3MDQ2 | Aldehyde dehydrogenase pseudogene | Mid |
| SULA_2758 | A0A0E3K1Q6 | Glutamate dehydrogenase | Start |
| SULA_2778 | A0A0E3JZ60 | Amidohydrolase | Mid |
| SULA_2847 | A0A0E3MC65 | ISC1439B family transposase | Mid |
| SULA_2865 | P95900 | L-lactate permease | End |
| SULA_03930 | A0A3G8ETZ4 | Hypothetical protein | Intergenic |
| SULA_03935 | N/A | Rhodonase domain protein (pseudogene) | |
| SULA_2891 | P95926 | Orf c01028 protein | |
| SULA_2892 | A0A0E3MDX3 | ISNCY family transposase | |

**Table S3. Proteins appearing in the Cren7 interactome and their abundance for each condition.**

| Protein Accession # | ORF | Overlap with Sso7D | Ref. # from Fig. 2 | Annotated Function | Abundance (fmol) | | |
|---|---|---|---|---|---|---|---|
| | | | | | L1 at pH 3 | L3 at pH 1 | L3 at ph 3 |
| A0A0E3JZK9 | SULA_0044 | | | Uncharacterized protein | 150±0 | 185±1 | 89±20 |
| A0A0E3GVT9 | SULA_0054 | | | Uncharacterized protein | 41±4 | 43±16 | 0±0 |
| A0A0E3GUA3 | SULA_0069 | | | Alkyl hydroperoxide reductase | 9±11 | 30±28 | 32±0 |
| A0A0E3K4P9 | SULA_0114 | X | | Cobalt-precorrin-3B C(17)-methyltransferase | 0±0 | 41±4 | 0±0 |
| A0A0E3M9B1 | SULA_0115 | X | | Precorrin-8X methylmutase | 0±0 | 304±10 | 0±0 |
| A0A0E3MCA7 | SULA_0162 | | | Succinate dehydrogenase | 90±13 | 109±22 | 29±12 |
| A0A0E3GVV9 | SULA_0165 | | | Succinate dehydrogenase | 0±6 | 0±36 | 1645±458 |
| A0A0E3MBQ7 | SULA_0179 | | | Translation initiation factor 1A | 15±4 | 11±9 | 11±6 |
| A0A0E3MEP5 | SULA_0182 | X | | Translation initiation factor 2 subunit beta | 21±30 | 18±7 | 15±25 |
| A0A0E3K6D7 | SULA_0215 | | | Thioredoxin reductase | 54±23 | 148±25 | 20±4 |
| A0A0E3MCK2 | SULA_0223 | X | | Heat-shock protein Hsp20 | 61±0 | 625±330 | 123±5 |
| A0A0E3M9H3 | SULA_0244 | | 27 | Putative Ruv-B like protien / TIP49 TBP-interacting protein | 136±24 | 225±60 | 15±3 |
| A0A0E3MG25 | SULA_0267 | | 18 | TrmB family transcriptional regulator | 9±8 | 19±28 | 40±19 |
| A0A0E3JSX5 | SULA_0268 | | 25 | DNA-binding protein | 238±43 | 281±11 | 172±42 |
| A0A0E3K4V7 | SULA_0288 | X | | DNA binding protein Cren8 | 37±4 | 50±16 | 45±14 |
| A0A0E3K740 | SULA_0302 | | | 3-ketoacyl-ACP reductase | 16±9 | 47±31 | 15±1 |
| A0A0E3KB67 | SULA_0379 | | | Zinc-containing ferredoxin | 2±3 | 23±5 | 23±14 |
| A0A0E3MG69 | SULA_0391 | | | 2-methylisocitrate lyase | 237±331 | 684±68 | 262±326 |
| A0A0E3K4Z9 | SULA_0392 | | | Histidine kinase | 12±0 | 51±54 | 44±37 |
| A0A0E3GTT8 | SULA_0433 | X | | Response regulator SirA | 7±2 | 19±1 | 14±1 |
| A0A0E3KAM8 | SULA_0436 | X | | Uncharacterized protein | 105±52 | 112±18 | 180±222 |
| A0A0E3M9L6 | SULA_0441 | | | xanthine dehydrogenase family protein subunit M | 26±1 | 57±2 | 24±1 |
| A0A0E3K6M1 | SULA_0443 | | | xanthine dehydrogenase family protein molybdopterin-binding subunit | 352±441 | 106±16 | 16±14 |
| A0A0E3K011 | SULA_0551 | | | Rubrerythrin | 136±80 | 137±5 | 21±39 |
| A0A0E3MF90 | SULA_0565 | | | Acyl-CoA dehydrogenase | 0±15 | 39±8 | 4±9 |
| A0A0E3MCJ0 | SULA_0566 | | | Electron transfer flavoprotein subunit alpha | 0±1 | 35±15 | 11±3 |
| A0A0E3MGH3 | SULA_0634 | | 13 | Putative transcription factor | 29±0 | 54±2 | 6±1 |
| A0A0E3KAJ7 | SULA_0658 | | | 3-hydroxypropionyl-CoA synthetase | 143±37 | 330±29 | 123±10 |
| A0A0E3MF59 | SULA_0662 | | | Uncharacterized protein | 57±34 | 66±27 | 231±24 |
| A0A0E3MD00 | SULA_0832 | | | Aldo/keto reductase | 211±50 | 499±36 | 26±13 |
| A0A0E3MDW0 | SULA_0884 | | | Dihydroxy-acid dehydratase | 49±32 | 115±7 | 27±9 |
| A0A0E3MGU8 | SULA_0968 | | | Mandelate racemase | 23±8 | 127±12 | 78±106 |
| A0A0E3K5J6 | SULA_1008 | | 12 | Winged helix-turn-helix transcriptional regulator | 33±13 | 64±4 | 34±1 |
| A0A0E3MDZ7 | SULA_1036 | | | Molybdenum cofactor biosynthesis protein MoaB | 13±1 | 57±50 | 8±2 |
| A0A0E3MD01 | SULA_1045 | | | Histidine kinase | 87±31 | 127±1 | 48±2 |
| A0A0E3K7F2 | SULA_1080 | | | Isopentenyl-diphosphate delta-isomerase | 20±1 | 45±16 | 23±14 |
| A0A0E3GUT2 | SULA_1086 | | | 30S ribosomal protein S9 | 135±13 | 158±12 | 83±31 |
| A0A0E3K7F5 | SULA_1087 | | | 50S ribosomal protein L13 | 34±10 | 55±1 | 40±7 |
| A0A0E3KB49 | SULA_1088 | X | | 50S ribosomal protein L18e | 66±9 | 85±3 | 47±2 |
| A0A0E3GWC1 | SULA_1089 | | 14 | DNA-directed RNA polymerase subunit Rpo3 | 28±4 | 44±2 | 15±2 |

| | | | | | | | |
|---|---|---|---|---|---|---|---|
| A0A0E3MIL5 | SULA_1090 | | | 30S ribosomal protein S11 | 63±6 | 254±17 | 63±11 |
| A0A0E3GUT3 | SULA_1091 | | | 30S ribosomal protein S4 | 81±11 | 125±31 | 55±21 |
| A0A0E3GT94 | SULA_1096 | | 26 | DNA primase DnaG | 196±188 | 497±55 | 25±19 |
| P95966 | SULA_1109 | | | Orf c04027 protein | 80±43 | 74±8 | 68±19 |
| A0A0E3KB68 | SULA_1119 | | | Aminotransferase | 173±164 | 204±139 | 51±1 |
| P95951 | SULA_1124 | | | Histidine kinase | 55±1 | 75±5 | 55±1 |
| A0A0E3K0D0 | SULA_1192 | | 11 | AAA ATPase | 34±20 | 61±30 | 12±4 |
| A0A0E3KB91 | SULA_1197 | | | Delta-aminolevulinic acid dehydratase | 81±1 | 150±66 | 51±17 |
| A0A0E3MIU3 | SULA_1216 | | 3 | S-adenosylmethionine synthase | 113±2 | 147±1 | 55±28 |
| A0A0E3KB97 | SULA_1217 | X | 21 | Sm ribonucleo | 3±7 | 41±39 | 21±0 |
| A0A0E3GWE3 | SULA_1222 | | | Uncharacterized protein | 25±0 | 138±44 | 41±22 |
| A0A0E3K0E4 | SULA_1224 | X | | Phosphoglucomutase | 0±3 | 127±86 | 21±0 |
| A0A0E3MFR0 | SULA_1237 | | | 30S ribosomal protein S7 | 56±24 | 237±220 | 79±0 |
| A0A0E3MD56 | SULA_1239 | | | 30S ribosomal protein S12 | 43±3 | 84±20 | 31±13 |
| A0A0E3GTB9 | SULA_1247 | | | Nucleoside diphosphate kinase | 2±0 | 35±7 | 39±4 |
| A0A0E3K207 | SULA_1249 | | | 30S ribosomal protein S28e | 28±2 | 41±4 | 22±0 |
| A0A0E3GTC2 | SULA_1265 | | | HIT family hydrolase | 0±0 | 11±15 | 23±3 |
| A0A0E3K8S0 | SULA_1293 | X | 7 | Sm ribonucleo | 149±18 | 160±12 | 60±9 |
| A0A0E3JXB9 | SULA_1311 | | 15 | DNA-directed RNA polymerase subunit Rpo11 | 27±25 | 74±34 | 0±0 |
| A0A0E3MIX4 | SULA_1314 | X | | 50S ribosomal protein L10e | 2574±123 | 2840±315 | 27±18 |
| A0A0E3K0I0 | SULA_1326 | | | Shikimate kinase | 92±42 | 57±23 | 28±23 |
| A0A0E3GX98 | SULA_1337 | | | Uncharacterized protein | 96±22 | 205±39 | 61±10 |
| A0A0E3JXD6 | SULA_1359 | | | 50S ribosomal protein L10 | 3636±97 | 4075±476 | 58±5 |
| A0A0E3GWG7 | SULA_1360 | X | | 50S ribosomal protein L1 | 40±4 | 69±6 | 58±0 |
| A0A0E3JVC2 | SULA_1361 | X | | 50S ribosomal protein L11 | 0±4 | 12±0 | 36±0 |
| A0A0E3GTE1 | SULA_1371 | | | 30S ribosomal protein S19e | 49±0 | 74±5 | 49±0 |
| A0A0E3MBX9 | SULA_1382 | X | | GTP cyclohydrolase 1 | 22±7 | 5±0 | 70±7 |
| A0A0E3MCQ0 | SULA_1415 | | | tRNA pseudouridine synthase B | 0±0 | 53±76 | 0±0 |
| A0A0E3K7T1 | SULA_1417 | | | 50S ribosomal protein L14e | 15±1 | 47±1 | 39±28 |
| A0A0E3MCQ4 | SULA_1430 | | 2 | Vitamin B12 Independent methionine synthase | 126±3 | 153±4 | 509±315 |
| A0A0E3MAB5 | SULA_1431 | | 1 | Vitamin B12 Dependent methionine synthase | 318±88 | 315±19 | 442±1 |
| A0A0E3GUZ1 | SULA_1432 | | | 30S ribosomal protein S15 | 146±1 | 182±1 | 37±9 |
| A0A0E3MFV7 | SULA_1434 | | | 30S ribosomal protein S6e | 51±8 | 76±1 | 49±5 |
| A0A0E3MBZ4 | SULA_1437 | | 22 | DNA-directed RNA polymerase subunit rpo4 | 1±0 | 2±1 | 139±17 |
| A0A0E3MBI9 | SULA_1443 | | | Nitrilase | 78±37 | 90±104 | 48±2 |
| A0A0E3MEA6 | SULA_1444 | | | 30S ribosomal protein S25 | 38±6 | 64±22 | 9±9 |
| A0A0E3GWI4 | SULA_1455 | | | 30S ribosomal protein S24e | 22±9 | 232±285 | 27±1 |
| A0A0E3MKK6 | SULA_1467 | | 9 | Transcription initiation factor IIB | 73±65 | 40±22 | 35±13 |
| A0A0E3K7W1 | SULA_1477 | | 32 | CRISPR-associated protein | 37±14 | 121±52 | 41±21 |
| A0A0E3MAC6 | SULA_1481 | | 16 | TrmB family transcriptional regulator | 14±4 | 19±2 | 18±6 |
| A0A0E3K900 | SULA_1483 | | 30 | Putative chromosome partitioning ATPase | 73±4 | 105±7 | 56±4 |
| A0A0E3MG44 | SULA_1531 | | 17 | AbrB family transcriptional regulator | 11±0 | 29±2 | 0±0 |
| A0A0E3GV26 | SULA_1634 | | | 3-hydroxy-3-methylglutaryl CoA synthase | 44±1 | 77±11 | 0±1 |
| Q9UWT8 | SULA_1635 | | | Acetyl CoA synthase | 0±0 | 87±54 | 0±5 |
| Q9UWT7 | SULA_1636 | X | 34 | DNA-binding protein | 14±13 | 29±5 | 8±2 |
| A0A0E3KCJ4 | SULA_1638 | X | 4 | Serine hydroxymethyltransferase | 79±7 | 1085±1091 | 115±91 |
| Q9UWT4 | SULA_1639 | | | Universal stress protein A | 0±0 | 49±13 | 0±0 |

| | | | | | | | |
|---|---|---|---|---|---|---|---|
| Q9UWV7 | SULA_1649 | | | Uncharacterized protein ORF-c21_046 | 29±21 | 52±7 | 30±7 |
| A0A0E3MJ87 | SULA_1658 | | | V-type ATP synthase alpha chain | 671±45 | 859±24 | 378±481 |
| A0A0E3GWL9 | SULA_1659 | | | V-type ATP synthase beta chain | 45±6 | 64±3 | 149±31 |
| A0A0E3MCW1 | SULA_1664 | X | | 30S ribosomal protein S26 | 20±4 | 30±24 | 14±0 |
| A0A0E3GTJ2 | SULA_1665 | | | Pyridoxal 5'-phosphate synthase subunit PdxS | 1236±272 | 734±609 | 370±460 |
| A0A0E3KCK7 | SULA_1666 | | | Pyridoxal 5'-phosphate synthase subunit PdxT | 99±20 | 123±1 | 309±56 |
| Q9UWZ9 | SULA_1697 | | 19 | Transcription factor S | 9±1 | 25±28 | 3±0 |
| A0A0E3GWN2 | SULA_1728 | | | Argininosuccinate synthase | 8±14 | 253±14 | 35±2 |
| Q9UX44 | SULA_1740 | | | Crotonase | 24±37 | 0±0 | 39±9 |
| A0A0E3GWN6 | SULA_1748 | | | Biotin--acetyl-CoA-carboxylase ligase | 143±20 | 120±67 | 47±28 |
| A0A0E3JXU6 | SULA_1782 | X | | 50S ribosomal protein L15 | 26±3 | 137±2 | 196±2 |
| A0A0E3KBZ7 | SULA_1783 | X | | 50S ribosomal protein L30 | 53±13 | 97±7 | 54±2 |
| A0A0E3GVR4 | SULA_1785 | | | 50S ribosomal protein L18 | 45±0 | 72±3 | 59±7 |
| A0A0E3KCR2 | SULA_1786 | | | 50S ribosomal protein L19e | 54±9 | 97±10 | 46±17 |
| A0A0E3MIX0 | SULA_1790 | X | | 30S ribosomal protein S14 type Z | 26±1 | 44±5 | 0±0 |
| A0A0E3GV55 | SULA_1792 | | | 30S ribosomal protein S4e | 69±0 | 86±6 | 54±7 |
| A0A0E3GTL6 | SULA_1794 | | | 50S ribosomal protein L14 | 19±1 | 45±14 | 11±12 |
| A0A0E3MEN5 | SULA_1795 | | | 30S ribosomal protein S17 | 51±0 | 27±0 | 83±9 |
| A0A0E3GV56 | SULA_1797 | X | | 50S ribosomal protein L29 | 38±0 | 56±3 | 44±0 |
| A0A0E3JUA0 | SULA_1800 | | | 30S ribosomal protein S19P | 34±15 | 49±11 | 30±10 |
| A0A0E3MG43 | SULA_1801 | | | 50S ribosomal protein L2 | 369±474 | 93±15 | 19±7 |
| A0A0E3GV57 | SULA_1802 | | | 50S ribosomal protein L23 | 50±36 | 94±9 | 50±3 |
| A0A0E3GWP6 | SULA_1803 | | | 50S ribosomal protein L4 | 67±1 | 148±18 | 31±24 |
| A0A0E3KC04 | SULA_1804 | | | 50S ribosomal protein L3 | 101±11 | 137±6 | 51±6 |
| A0A0E3KCS3 | SULA_1807 | | | 3-isopropylmalate dehydrogenase | 60±0 | 73±7 | 38±22 |
| A0A0E3KC07 | SULA_1811 | | | Elongation factor 2 | 318±353 | 582±413 | 170±23 |
| A0A0E3K0Y4 | SULA_1816 | X | | 50S ribosomal protein L37Ae | 80±91 | 97±108 | 9±2 |
| A0A0E3MHT5 | SULA_1817 | | | Exosome complex component Rrp42 | 18±1 | 58±0 | 25±19 |
| A0A0E3GWQ0 | SULA_1825 | | | 50S ribosomal protein L15e | 43±1 | 82±5 | 30±0 |
| A0A0E3MHT9 | SULA_1828 | | | 30S ribosomal protein S3Ae | 233±18 | 118±16 | 234±17 |
| Q9UXD9 | SULA_1833 | | 6 | DNA-directed RNA polymerase subunit Rpo7 | 1641±245 | 1868±369 | 36±1 |
| A0A0E3MGJ6 | SULA_1834 | | | 50S ribosomal protein L21e | 20±0 | 74±3 | 14±1 |
| Q9UXE9 | SULA_1851 | | | Uncharacterized protein ORF-c20_053 | 51±3 | 74±4 | 65±13 |
| Q9UXF0 | SULA_1852 | | | Uncharacterized protein ORF-c20_054 | 325±295 | 182±1 | 181±7 |
| A0A0E3K8E7 | SULA_1857 | | 35 | Replication factor C small subunit | 2±7 | 27±13 | 23±18 |
| A0A0E3MGL5 | SULA_1872 | X | 31 | Putative DNA binding protein | 55±67 | 44±52 | 0±0 |
| A0A0E3MD43 | SULA_1882 | | | Thermosome subunit | 170±56 | 748±26 | 226±78 |
| A0A0E3KC25 | SULA_1908 | | | Aminotransferase | 22±36 | 102±14 | 123±61 |
| A0A0E3K9H1 | SULA_1925 | | | phosphopyruvate hydratase | 13±1 | 67±24 | 9±7 |
| A0A0E3GV80 | SULA_1936 | | | Fe-S cluster assembly ATPase SufC | 33±4 | 85±8 | 0±12 |
| A0A0E3MH21 | SULA_1941 | | | Vitamin B12-dependent ribonucleotide reductase | 931±122 | 2633±61 | 731±91 |
| A0A0E3JUC9 | SULA_1948 | | 8 | Fibrillarin-like rRNA/tRNA 2'-O-methyltransferase | 84±10 | 57±0 | 45±14 |
| A0A0E3GTP4 | SULA_1957 | X | | Uncharacterized protein | 69±84 | 92±90 | 25±2 |
| A0A0E3GXA5 | SULA_1959 | | 10 | TATA-box-binding protein | 46±66 | 20±10 | 55±13 |
| A0A0E3MBJ8 | SULA_1972 | | 24 | DNA/RNA-binding protein Alba | 866±12 | 1484±4 | 871±132 |
| A0A0E3K8J6 | SULA_1977 | | | tRNA-splicing ligase RtcB | 10±15 | 69±73 | 18±7 |
| A0A0E3K654 | SULA_1979 | | 28 | Type 2 DNA topoisomerase 6 subunit B | 103±127 | 192±107 | 38±0 |

| | | | | | | | |
|---|---|---|---|---|---|---|---|
| A0A0E3MJJ4 | SULA_1981 | X | | Translation initiation factor 5A | 2±4 | 159±13 | 232±32 |
| A0A0E3MDS4 | SULA_1984 | | | 3-hydroxyisobutyrate dehydrogenase | 5±2 | 83±90 | 8±5 |
| A0A0E3GV89 | SULA_1986 | | 23 | Chromatin protein Cren7 | 13867±11 | 18918±1 | 15708±46 |
| A0A0E3GVA0 | SULA_2054 | X | | 50S ribosomal protein L44e | 11±0 | 20±2 | 16±3 |
| A0A0E3GTR1 | SULA_2056 | | | Translation initiation factor 2 subunit alpha | 20±14 | 68±5 | 32±0 |
| A0A0E3MGA9 | SULA_2067 | X | 5 | Probable glycine cleavage system H protein | 11±6 | 8±1 | 52±5 |
| A0A0E3MGS9 | SULA_2074 | | | Peptidase | 19±11 | 55±10 | 30±4 |
| A0A0E3MCH5 | SULA_2085 | X | | Fumarate hydratase class II | 24±12 | 436±613 | 0±6 |
| A0A0E3GTR8 | SULA_2100 | | | Aconitate hydratase | 135±41 | 213±1 | 158±45 |
| A0A0E3JUG3 | SULA_2102 | | | Uncharacterized protein | 84±130 | 164±189 | 15±5 |
| A0A0E3JY62 | SULA_2106 | | 20 | TrmB family transcriptional regulator | 8±7 | 11±2 | 165±49 |
| A0A0E3MGA7 | SULA_2116 | | | CoA-binding protein | 0±0 | 50±2 | 0±0 |
| A0A0E3JUH1 | SULA_2128 | | | Peroxiredoxin | 0±1 | 21±1 | 11±10 |
| A0A0E3K9S1 | SULA_2129 | | | Response regulator SirA | 94±0 | 91±16 | 75±43 |
| A0A0E3JY87 | SULA_2144 | | | Tryptophan synthase beta chain | 0±0 | 115±48 | 57±5 |
| A0A0E3MCJ2 | SULA_2151 | | | Peptidase U62 | 28±47 | 120±87 | 78±21 |
| A0A0E3KCF1 | SULA_2154 | | | MaoC family dehydratase | 0±0 | 8±12 | 31±0 |
| A0A0E3KCH6 | SULA_2224 | | | Uncharacterized protein | 7±1 | 64±36 | 84±18 |
| A0A0E3KDL8 | SULA_2657 | | | Uncharacterized protein | 9±13 | 172±8 | 97±79 |
| A0A0E3MIM1 | SULA_2718 | | 29 | DNA-binding protein Sso7d | 86±121 | 131±185 | 0±0 |
| A0A0E3JZ60 | SULA_2778 | | | Amidopeptidase | 52±14 | 42±17 | 69±29 |
| A0A0E3GU31 | SULA_2802 | X | | Acetyl-CoA acetyltransferase | 60±24 | 78±2 | 59±2 |
| A0A0E3MGV7 | SULA_2804 | | 33 | DNA-binding protein | 23±22 | 78±28 | 1±1 |

**Table S4. Proteins appearing in the Sso7D interactome.**

| Protein Accession # | ORF | Overlap with Cren7 | Ref. # from Fig. 2 | Annotated Function | Abundance (fmol) | | |
|---|---|---|---|---|---|---|---|
| | | | | | L1 at pH 3 | L3 at pH 1 | L3 at pH 3 |
| A0A0E3K074 | SULA_0896 | | | Fumarylacetoacetate hydrolase | 0±0 | 18±1 | 25±19 |
| A0A0E3GWG7 | SULA_1360 | X | | 50S ribosomal protein L1 | 33±2 | 82±58 | 164±53 |
| A0A0E3JY74 | SULA_2113 | | 3 | TrmB family transcriptional regulator | 208±6 | 160±8 | 223±91 |
| A0A0E3KB49 | SULA_1088 | X | | 50S ribosomal protein L18e | 25±12 | 14±3 | 32±0 |
| A0A0E3K0C2 | SULA_1169 | | | [LysW]-L-2-aminoadipate/[LysW]-L-glutamate phosphate reductase | 82±5 | 67±4 | 50±12 |
| A0A0E3MGA9 | SULA_2067 | X | 2 | Probable glycine cleavage system H protein | 3±4 | 2±4 | 3±2 |
| A0A0E3GU31 | SULA_2802 | X | | Acetyl-CoA acetyltransferase | 68±76 | 27±8 | 26±2 |
| A0A0E3MD09 | SULA_1738 | | | Adenosine monophosphate-protein transferase | 52±74 | 13±19 | 38±58 |
| A0A0E3JTR1 | SULA_1220 | | | D-arabino 3-hexulose 6-phosphate aldehyde lyase | 21±0 | 4±2 | 5±2 |
| A0A0E3KA78 | SULA_0352 | | 8 | DNA-binding protein Sso7D | 2055±1 | 1575±8 | 3427±9 |
| A0A0E3K5Z4 | SULA_1663 | | | Proline--tRNA ligase | 217±19 | 215±5 | 177±101 |
| A0A0E3MIX4 | SULA_1314 | X | | 50S ribosomal protein L10e | 126±19 | 213±62 | 203±141 |
| A0A0E3M9Q5 | SULA_0625 | | | Electron transfer flavoprotein subunit alpha | 162±10 | 133±1 | 178±48 |
| A0A0E3K6C7 | SULA_2265 | | | Peptide ABC transporter ATP-binding protein | 231±17 | 124±91 | 0±0 |
| A0A0E3KCJ4 | SULA_1638 | X | 1 | Serine hydroxymethyltransferase | 90±22 | 61±54 | 141±186 |
| A0A0E3JYC2 | SULA_2246 | | | Citryl-CoA lyase | 86±28 | 83±1 | 137±16 |
| A0A0E3KAM8 | SULA_0436 | X | | Uncharacterized protein | 17±9 | 74±86 | 168±158 |
| A0A0E3K5T7 | SULA_1410 | | | Glycosylated S-layer protein, SlaB | 84±16 | 24±35 | 133±37 |
| A0A0E3MCK2 | SULA_0223 | X | | Heat-shock protein Hsp20 | 29±28 | 53±14 | 245±311 |
| A0A0E3MAT5 | SULA_0013 | | | Membrane protease subunit, stomatin/prohibitin-like protein | 64±22 | 71±0 | 83±29 |
| A0A0E3KBZ7 | SULA_1783 | X | | 50S ribosomal protein L30 | 57±29 | 22±2 | 67±52 |
| A0A0E3KD23 | SULA_2018 | | | Pyruvate dehydrogenase | 0±0 | 132±17 | 13±14 |
| A0A0E3K0E4 | SULA_1224 | X | | Phosphoglucomutase | 89±159 | 0±7 | 104±139 |
| A0A0E3MCH5 | SULA_2085 | X | | Fumarate hydratase class II | 8±9 | 0±2 | 142±266 |
| A0A0E3JXU6 | SULA_1782 | X | | 50S ribosomal protein L15 | 33±12 | 46±2 | 46±24 |
| A0A0E3GUE2 | SULA_0285 | | | Succinate--CoA ligase [ADP-forming] subunit alpha | 38±0 | 38±11 | 45±23 |
| A0A0E3KD16 | SULA_1998 | | | Alpha-amylase | 29±28 | 62±6 | 14±4 |
| A0A0E3MJJ4 | SULA_1981 | X | | Translation initiation factor 5A | 5±16 | 6±15 | 113±7 |
| A0A0E3MFS1 | SULA_0006 | | | Uncharacterized protein | 30±1 | 37±10 | 38±4 |
| A0A0E3JXC8 | SULA_1339 | | | Uncharacterized protein | 22±21 | 6±1 | 124±173 |
| A0A0E3MD09 | SULA_1738 | | | Adenosine monophosphate-protein transferase | 52±74 | 13±19 | 38±58 |
| A0A0E3GUF7 | SULA_0374 | | | Uncharacterized protein | 16±16 | 1±0 | 64±51 |
| A0A0E3MFG4 | SULA_0762 | | | Putative methylthioribose-1-phosphate isomerase | 20±6 | 14±1 | 34±66 |
| A0A0E3JW49 | SULA_0383 | | | Uncharacterized protein | 0±0 | 59±33 | 0±0 |
| A0A0E3MFW9 | SULA_0117 | | | MBL fold metallo-hydrolase | 8±10 | 50±41 | 0±0 |
| A0A0E3MIX0 | SULA_1790 | X | | 30S ribosomal protein S14 type Z | 26±3 | 29±1 | 0±0 |
| A0A0E3K517 | SULA_0454 | | 4 | AsnC family transcriptional regulator | 19±0 | 16±0 | 26±1 |
| A0A0E3GVA0 | SULA_2054 | X | | 50S ribosomal protein L44e | 20±7 | 29±1 | 10±12 |

| | | | | | | | |
|---|---|---|---|---|---|---|---|
| A0A0E3ME83 | SULA_2194 | | | Phosphohistidine phosphatase | 7±0 | 10±0 | 56±73 |
| A0A0E3GWP4 | SULA_1793 | | | 50S ribosomal protein L24 | 31±11 | 22±6 | 0±12 |
| A0A0E3KB91 | SULA_1197 | | | Delta-aminolevulinic acid dehydratase | 16±5 | 22±0 | 5±5 |
| A0A0E3MCW1 | SULA_1664 | X | | 30S ribosomal protein S26 | 20±7 | 25±2 | 0±1 |
| A0A0E3GTP4 | SULA_1957 | X | | Uncharacterized protein | 20±1 | 16±1 | 11±12 |
| A0A0E3GV56 | SULA_1797 | X | | 50S ribosomal protein L29 | 13±9 | 6±0 | 26±21 |
| A0A0E3K8S0 | SULA_1293 | X | 5 | Sm ribonucleo | 5±3 | 22±17 | 16±20 |
| A0A0E3GUG7 | SULA_0429 | | | Acetyl-CoA acetyltransferase | 3±2 | 30±65 | 2±1 |
| A0A0E3MEY1 | SULA_0107 | | | Acyl-CoA Thioesterase | 29±31 | 3±4 | 0±0 |
| A0A0E3MAW3 | SULA_2104 | | | Uncharacterized protein | 20±11 | 8±12 | 9±0 |
| A0A0E3K0Y4 | SULA_1816 | X | | 50S ribosomal protein L37Ae | 10±0 | 13±0 | 9±7 |
| A0A0E3ME51 | SULA_1248 | | | 50S ribosomal protein L24e | 16±0 | 10±0 | 0±0 |
| A0A0E3MEP5 | SULA_0182 | X | | Translation initiation factor 2 subunit beta | 3±0 | 3±0 | 25±11 |
| A0A0E3GTT8 | SULA_0433 | X | | Response regulator SirA | 0±0 | 0±0 | 51±67 |
| A0A0E3M9J2 | SULA_0401 | | | alanine--glyoxylate aminotransferase family protein | 10±2 | 2±4 | 19±27 |
| A0A0E3MHE0 | SULA_0139 | | | Putative cobyrinic acid a,c-diamide synthase | 9±0 | 26±3 | 0±10 |
| A0A0E3K4V7 | SULA_0288 | X | | DNA binding protein | 5±5 | 2±0 | 15±0 |
| A0A0E3MKX2 | SULA_1342 | | | NADH dehydrogenase | 0±14 | 6±11 | 20±7 |
| A0A0E3KB97 | SULA_1217 | X | 6 | Sm ribonucleo | 14±3 | 7±11 | 0±0 |
| A0A0E3K597 | SULA_0671 | | | Alcohol dehydrogenase | 0±0 | 16±1 | 0±0 |
| A0A0E3JVC2 | SULA_1361 | X | | 50S ribosomal protein L11 | 14±2 | 3±7 | 4±7 |
| A0A0E3MAE0 | SULA_1440 | | | 2,3-bisphosphoglycerate-independent phosphoglycerate mutase | 2±3 | 13±1 | 5±15 |
| A0A0E3M9B1 | SULA_0115 | X | | Precorrin-8X methylmutase | 0±0 | 14±5 | 0±0 |
| A0A0E3MGL5 | SULA_1872 | X | | UPF0148 protein SULA_1872 | 5±4 | 5±6 | 0±0 |
| A0A0E3MGH3 | SULA_063 | | | Uncharacterized protein | 23±2 | 7±5 | 0±19 |
| Q9UWT7 | SULA_1636 | X | 9 | DNA-binding protein | 0±0 | 9±4 | 0±1 |
| A0A0E3MHM4 | SULA_1661 | | | ATPase | 3±0 | 0±0 | 8±5 |
| A0A0E3MEB1 | SULA_0095 | | | Uncharacterized protein | 3±3 | 5±1 | 0±0 |
| A0A0E3K4P9 | SULA_0114 | X | | Cobalt-precorrin-3B C(17)-methyltransferase | 0±0 | 8±0 | 0±0 |
| A0A0E3JUF4 | SULA_2073 | | 7 | LysR family transcriptional regulator | 2±1 | 4±0 | 2±5 |
| A0A0E3K074 | SULA_0896 | | | Fumarylacetoacetate hydrolase | 0±0 | 18±1 | 25±19 |
| A0A0E3GWG7 | SULA_1360 | X | | 50S ribosomal protein L1 | 33±2 | 82±58 | 164±53 |

**Table S5. Sequencing adaptors used in this work**.

| Primer | Sequence (5'-3') | Reference |
|---|---|---|
| Single read adapter A | NNNNNNAGATCGGAAGAGCTCGTATGCCGTCTTCTGCTTG | (6) |
| Single read adapter B | ACACTCTTTCCCTACACGACGCTCTTCCGATCTTGCGTTT | (6) |
| NNNNNN = 6mer barcode used in each sample | | |

**Table S6. Summary of ChIPseq library assemblies.**

| Sample | Reads | Aligned reads | Aligned reads % | Reads failed to align | Reads failed to align (%) | Coverage (reads/bp) |
|---|---|---|---|---|---|---|
| Cren7 SARC pH1 - 1 | 18734126 | 17482780 | 93% | 1251346 | 7% | 6.65 |
| Cren7 SARC pH1 - 2 | 13802814 | 12958345 | 94% | 844469 | 6% | 4.93 |
| Cren7 SARC pH1 - 3 | 16050826 | 15096445 | 94% | 954381 | 6% | 5.74 |
| Cren7 SARC pH3 - 1 | 17445426 | 16350824 | 94% | 1094602 | 6% | 6.22 |
| Cren7 SARC pH3 - 2 | 17664916 | 17105086 | 97% | 559830 | 3% | 6.50 |
| Cren7 SARC pH3 - 3 | 16789800 | 16147673 | 96% | 642127 | 4% | 6.14 |
| Cren7 WT pH3 -1 | 8364344 | 7517927 | 90% | 846417 | 10% | 2.86 |
| Cren7 WT pH3 - 2 | 17664916 | 17105086 | 97% | 559830 | 3% | 6.50 |
| Cren7 WT pH3 - 3 | 16542076 | 15772637 | 95% | 769439 | 5% | 6.00 |
| Sso7D SARC pH1 - 1 | 18310976 | 10983217 | 60% | 7327759 | 40% | 4.18 |
| Sso7D SARC pH1 - 2 | 23233494 | 13906203 | 60% | 9327291 | 40% | 5.29 |
| Sso7D SARC pH1 - 3 | 23233494 | 13906203 | 60% | 9327291 | 40% | 5.29 |
| Sso7D SARC pH3 - 1 | 26061184 | 15334354 | 59% | 10726830 | 41% | 5.83 |
| Sso7D SARC pH3 - 2 | 21041716 | 12983477 | 62% | 8058239 | 38% | 4.94 |
| Sso7D SARC pH3 - 3 | 22492798 | 13901728 | 62% | 8591070 | 38% | 5.29 |
| Sso7D WT pH3 -1 | 5065464 | 3844849 | 76% | 1220615 | 24% | 1.46 |
| Sso7D WT pH3 - 2 | 23203872 | 13974963 | 60% | 9228909 | 40% | 5.31 |
| Sso7D WT pH3 - 3 | 15871466 | 9771231 | 62% | 6100235 | 38% | 3.72 |

### References

1. Allen, M. B. (1959) Studies with Cyanidium caldarium, an anomalously pigmented chlorophyte. *Arch Mikrobiol* **32**, 270-277
2. Rolfsmeier, M., and Blum, P. (1995) Purification and characterization of a maltase from the extremely thermophilic crenarchaeote *Sulfolobus solfataricus*. *J Bacteriol* **177**, 482-485
3. Worthington, P., Blum, P., Perez-Pomares, F., and Elthon, T. (2003) Large-scale cultivation of acidophilic hyperthermophiles for recovery of secreted proteins. *Appl Environ Microbiol* **69**, 252-257
4. Brock, T. D., Brock, K. M., Belly, R. T., and Weiss, R. L. (1972) *Sulfolobus*: a new genus of sulfur-oxidizing bacteria living at low pH and high temperature. *Arch Mikrobiol* **84**, 54-68
5. Robinson, N. P., Dionne, I., Lundgren, M., Marsh, V. L., Bernander, R., and Bell, S. D. (2004) Identification of two origins of replication in the single chromosome of the archaeon Sulfolobus solfataricus. *Cell* **116**, 25-38
6. Rudrappa, D., Yao, A. I., White, D., Pavlik, B. J., Singh, R., Facciotti, M. T., and Blum, P. (2015) Identification of an archaeal mercury regulon by chromatin immunoprecipitation. *Microbiology* **161**, 2423-2433
7. Langmead, B., and Salzberg, S. L. (2012) Fast gapped-read alignment with Bowtie 2. *Nat Methods* **9**, 357-359
8. Li, H. (2011) A statistical framework for SNP calling, mutation discovery, association mapping and population genetical parameter estimation from sequencing data. *Bioinformatics* **27**, 2987-2993
9. Ramirez, F., Dundar, F., Diehl, S., Gruning, B. A., and Manke, T. (2014) deepTools: a flexible platform for exploring deep-sequencing data. *Nucleic Acids Res* **42**, W187-191
10. Arrigoni, L., Richter, A. S., Betancourt, E., Bruder, K., Diehl, S., Manke, T., and Bonisch, U. (2016) Standardizing chromatin research: a simple and universal method for ChIP-seq. *Nucleic Acids Res* **44**, e67
11. Turcan, S., Makarov, V., Taranda, J., Wang, Y., Fabius, A. W. M., Wu, W., Zheng, Y., El-Amine, N., Haddock, S., Nanjangud, G., LeKaye, H. C., Brennan, C., Cross, J., Huse, J. T., Kelleher, N. L., Osten, P., Thompson, C. B., and Chan, T. A. (2018) Mutant-IDH1-dependent chromatin state reprogramming, reversibility, and persistence. *Nat Genet* **50**, 62-72
12. Freese, N. H., Norris, D. C., and Loraine, A. E. (2016) Integrated genome browser: visual analytics platform for genomics. *Bioinformatics* **32**, 2089-2095
13. Thorvaldsdottir, H., Robinson, J. T., and Mesirov, J. P. (2013) Integrative Genomics Viewer (IGV): high-performance genomics data visualization and exploration. *Brief Bioinform* **14**, 178-192
14. Bailey, T. L., Boden, M., Buske, F. A., Frith, M., Grant, C. E., Clementi, L., Ren, J., Li, W. W., and Noble, W. S. (2009) MEME SUITE: tools for motif discovery and searching. *Nucleic Acids Res* **37**, W202-208
15. Payne, S., McCarthy, S., Johnson, T., North, E., and Blum, P. (2018) Nonmutational mechanism of inheritance in the Archaeon Sulfolobus solfataricus. *Proc Natl Acad Sci U S A* **115**, 12271-12276
16. Li, K. W., Chen, N., Klemmer, P., Koopmans, F., Karupothula, R., and Smit, A. B. (2012) Identifying true protein complex constituents in interaction proteomics: the example of the DMXL2 protein complex. *Proteomics* **12**, 2428-2432

17. Altschul, S. F., Madden, T. L., Schaffer, A. A., Zhang, J., Zhang, Z., Miller, W., and Lipman, D. J. (1997) Gapped BLAST and PSI-BLAST: a new generation of protein database search programs. *Nucleic Acids Res* **25**, 3389-3402
18. Marchler-Bauer, A., Anderson, J. B., Derbyshire, M. K., DeWeese-Scott, C., Gonzales, N. R., Gwadz, M., Hao, L., He, S., Hurwitz, D. I., Jackson, J. D., Ke, Z., Krylov, D., Lanczycki, C. J., Liebert, C. A., Liu, C., Lu, F., Lu, S., Marchler, G. H., Mullokandov, M., Song, J. S., Thanki, N., Yamashita, R. A., Yin, J. J., Zhang, D., and Bryant, S. H. (2007) CDD: a conserved domain database for interactive domain family analysis. *Nucleic Acids Res* **35**, D237-240
19. Ogata, H., Goto, S., Sato, K., Fujibuchi, W., Bono, H., and Kanehisa, M. (1999) KEGG: Kyoto Encyclopedia of Genes and Genomes. *Nucleic Acids Res* **27**, 29-34
20. Struys, E. A., Jansen, E. E., de Meer, K., and Jakobs, C. (2000) Determination of *S*-adenosylmethionine and *S*-adenosylhomocysteine in plasma and cerebrospinal fluid by stable-isotope dilution tandem mass spectrometry. *Clin Chem* **46**, 1650-1656
21. Luippold, G., Delabar, U., Kloor, D., and Muhlbauer, B. (1999) Simultaneous determination of adenosine, S-adenosylhomocysteine and S-adenosylmethionine in biological samples using solid-phase extraction and high-performance liquid chromatography (vol 724, pg 231, 1999). *J Chromatogr B* **729**, 379-379
22. Stabler, S. P., and Allen, R. H. (2004) Quantification of serum and urinary *S*-adenosylmethionine and *S*-adenosylhomocysteine by stable-isotope-dilution liquid chromatography-mass spectrometry. *Clin Chem* **50**, 365-372
